# Predictive and instructive cerebellar encoding of dopamine reward drives motivated behavior

**DOI:** 10.64898/2026.02.02.703324

**Authors:** Benjamin A. Filio, Amma Otchere, Subhiksha Srinivasan, Srijan Thota, Luke Drake, Lizmaylin Ramos, Philipp Maurus, Mark J. Wagner

**Affiliations:** National Institute of Neurological Disorders & Stroke, National Institutes of Health, Bethesda, MD 20894, USA; Department of Neuroscience, Brown University, Providence, RI 02906, USA

## Abstract

Learning motivated behaviors requires both anticipating rewards and reinforcing actions that yield them. Although cerebellar activity encodes natural rewards like water and food, it also coordinates physical movements like eating and drinking. To disentangle reward from consummatory movements, we trained mice to push for delayed dopamine rewards delivered directly into the brain. Here we show that cerebellar input streams use both predictive and instructive codes for dopamine reward. Two-photon imaging revealed that many cerebellar granule cells (GrCs) predictively encoded dopamine rewards with sustained activity that “stretched” to match 1- or 2-s delay intervals before terminating upon reward receipt. By contrast, most cerebellar climbing fibers (CFs) spiked just after dopamine delivery. In mice also trained with water rewards, encoding strength for dopamine matched or exceeded that for water. Both cell types contributed causally: chronic GrC inhibition disrupted self-stimulation learning, and CF self-stimulation “rewards” drove moderate operant learning in naive animals. Thus, cerebellar encoding of dopamine reward helps drive motivated behavior, suggesting deeper cerebellar integration in brain reward prediction networks.

## Main

Learning which actions yield desired outcomes involves reinforcing rewarded behaviors^1,2^—an important brain-wide process with key contributions from nigrostriatal circuits in which midbrain dopamine (DA) reward signals drive downstream associative learning in the striatum^3^. By contrast, the cerebellum has long been known to contribute to learning predictions^4,5^, such as near-simultaneous associations vital for sensorimotor functions^6,7^. Recent cerebellar studies have broadened the concept of prediction learning to include, among other things, “reward prediction.” In thirsty animals seeking water, various cerebellar neurons convey signals that could be useful for predicting water reward occurrence, estimating its timing, or associating it with actions or events^8–10^. On the other hand, extensive motor programs are required to “consume” physical rewards, like drinking water, eating food, and social interaction—a major confound given the cerebellum’s critical role in refining physical movements. Thus, although GrCs exhibit a variety of activity patterns predictive of water reward^11^, these signals might serve mainly to drive anticipatory licking or orofacial movements needed for water consumption. Similarly, although CFs convey heterogeneous reinforcement signals^12–14^, with transient spiking at water delivery or predictive cues or actions, these signals might instead help reinforce or refine drinking movements. Similar caveats confound cerebellar contributions to other ethological physical rewards like food^15^ and socialization^16^. Studies have also found that cerebellar Purkinje cells^11,17^ and cerebellar nuclear neurons^18^ convey reward- predicting signals, but these too employed rewards requiring physical consumption.

Together, these results raise a critical question: does cerebellar activity signal reward per se, or only when action is required to sense or consume it? If cerebellar circuits predictively encode rewards that do not need physical consumption, they may be more deeply integrated with broader brain reward prediction networks and their associated disorders like compulsive reward-seeking and substance use. To adjudicate these possibilities, we trained mice to push a robotic handle for delayed self-stimulation dopamine rewards that are delivered directly into the brain and thus passively consumed. Via two-photon imaging of both cerebellar input streams, GrCs and CFs, we found both predictive and instructive encoding of dopamine rewards. Numerous GrCs exhibited sustained anticipatory activity that was quenched by dopamine delivery, linking the action to the expected reward time. This activity temporally scaled to match 1- or 2-s delays. Dopamine delivery triggered widespread CF spiking. These patterns generalized between water and dopamine rewards, and thus likely encoded abstract features of rewarded behavior. Selectively perturbing delay-period GrC activity prevented normal learning of DA-motivated behavior. Conversely, providing only delayed optogenetic CF ‘rewards’ also drove naïve mice to push at moderate rates. Together, these data demonstrate widespread cerebellar encoding of DA reward and its causal contribution to learning and executing motivated behaviors, broadening possible cerebellar contributions to reward-driven behavior.

### Push-for-delayed-self-stimulation tasks compatible with two-photon cerebellar imaging

We sought an operant task where animals associate an action with a reward, with several features. First, animals should receive rewards without needing to “consume” them via physical movement—versus, e.g., water, which animals must drink and often begin licking for beforehand. This helps avoid conflating neural representations of reward with those of consumption movements. Second, we temporally delayed the rewards so that transient neural representations of the operant action did not “mix” with later reward prediction signals. Third, to study GrC–CF dynamics, the paradigm would be surgically, genetically, and optically compatible with our two-photon cerebellar imaging strategy^11^. We thus designed a head-fixed push-for-delayed-self-stimulation task (**Fig. 1a**). We focused on two complementary self-stimulation rewards: (1) optogenetic activation of DA neurons in the ventral tegmental area^19^ (VTA) using ChRimson (**Fig. 1b,c; Video S1**) and (2) electrical activation of the medial forebrain bundle^20^ (MFB) (**Fig. 1d,e, Video S2**) (both left side only, contralateral to the reaching forelimb). VTA activation benefited from selectively targeting DA populations that signal rewarding outcomes^21^ (albeit heterogeneously^22^), but suffered from high viral-genetic and optical complexity (see **Imaging** below). By contrast, MFB electrical activation was simpler but less specific than VTA activation: it drives motivation through multiple dopaminergic^23,24^ and non-dopaminergic pathways^25^. To obtain robust results, we studied both paradigms in parallel (MFB: 19 mice; VTA: 8 mice). Since findings were broadly consistent for both reward types, we hereafter write “DA reward” when referring to both.

**Figure 1.**
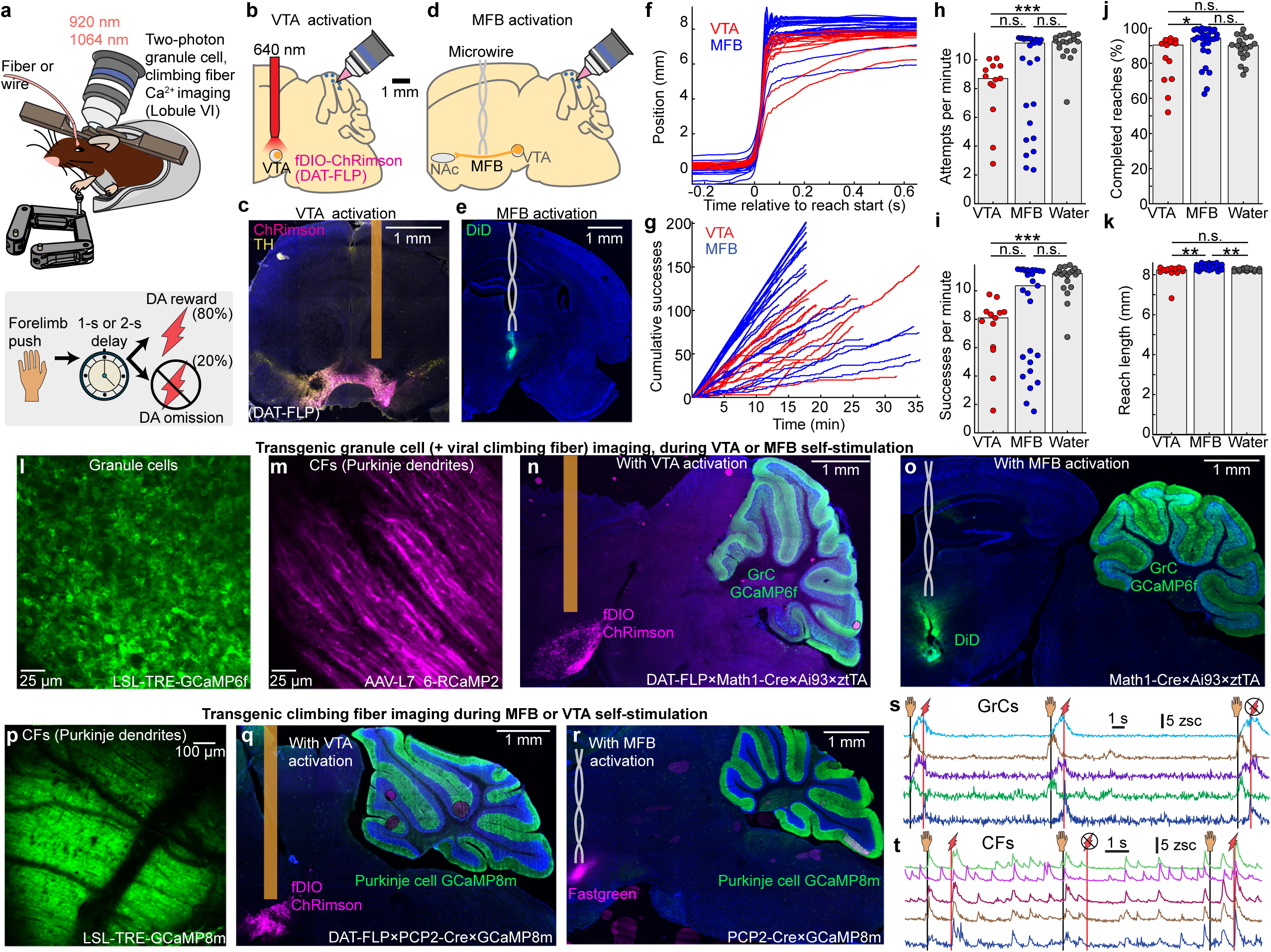
Push-for-delayed-self-stimulation tasks and simultaneous two-photon cerebellar imaging. **a-k**, Behavior. **a,** Task schematic. Head-fixed mice self-initiated right forelimb pushes of a robotic handle (maximum 8 mm extent) for delayed self-stimulation rewards. **b,c, VTA dopamine reward** (n=8 mice). Schematic (**b**) and histology (**c**) of optogenetic activation via FLP-dependent ChRimson in DAT-FLP mice (2- or 5-ms pulses at 50 Hz for 250 or 500 ms, 10– 15 mW at 640 nm). **d,e, Medial forebrain bundle (MFB) stimulation reward** (n=19 mice). Schematic (**d**) and histology (**e**) of electrical activation via microwire (5-ms 5-V pulses at 100 Hz for 250 ms). **f,** Handle position averaged across trials and sessions (sessions/mice: 13/8 VTA and 27/19 MFB). **g,** Cumulative successful reaches over time for each session. **h**–**k,** Quantification of expert behavior (after 5-7 training sessions). Reach attempts (**h**) and successes (**i**), computed as 1/median inter-trial interval (ITI). **j,** Completion percentage. **k,** Length of successful reaches. Points are session medians. Bars show medians and dots show sessions. (Bonferroni-corrected two-sided Mann-Whitney U-test p values VTA v MFB / MFB v Water / VTA v Water: **h,** 0.2 / 0.2 / 4x10^-5^; **i**, 0.3 / 0.2 / 5x10^-5^; **j,** 0.02 / 0.3 / 1; **k,** 0.007 / 0.002 / 1. Water *n*=19 sessions/15 mice). Learning metrics in **Extended Data Fig. 1**. **l-t**, Two-photon imaging during behavior. **l,** *In vivo* image of GrCs via Math1-Cre×LSL-tTA×LSL-TRE-GCaMP6f (20 mice). **m,** Purkinje dendrites (reporting CF activity) expressing viral R-CaMP2 in Math1-Cre×tTA×GCaMP6f mice (n=5). **n,o,** Histology for imaging GrCs during VTA activation (**n,** DAT-FLP×Math1-Cre×ztTA×GCaMP6f with fDIO-ChRimson in VTA, n=6) or MFB stimulation (**o**, DiD staining; n=14). **p,** *In vivo* two-photon image of transgenic GCaMP8m in Purkinje dendrites (PCP2-Cre×LSL-GCaMP8m; 6 mice). **q,r,** Histology for transgenic Purkinje dendrite imaging during VTA DA stimulation (**q**, DAT-FLP×PCP2-Cre×GCaMP8m; 2 mice) or MFB stimulation (PCP2-Cre×GCaMP8m mice; 4 mice). **s,t,** Fluorescence traces over 3 trials from 5 example GrCs (**s**) and CFs (**t**). Icons indicate push and reward/omission times.

We trained animals to self-initiate right forelimb pushes of a robotic manipulandum^26^ (constrained to linear motion, maximum extent: 8-mm; **Methods**). Movements >6 mm in length were rewarded 1 s after completion with self-stimulation (VTA: 2- or 5-ms pulses at 50 Hz for 250 ms [1 mouse] or 500 ms [7 mice], 10–15 mW of 640 nm laser power; MFB: 5 ms, 5 V pulses at 100 Hz for 250 ms). After another 2-s delay, the handle began automatically returning to the start position over the next 2 s before animals were able to initiate the next trial. Training lasted 5–7 days. Expert mice typically made 100 or more successful reaches in each session (**Fig. 1f,g**), making 8.7±0.5 and 11±0.9 attempts per minute and 8.1±0.6 and 10.4±1.5 successful reaches per minute, yielding overall completion rates of 90±6% and 94±1%, for VTA and MFB rewards respectively (**Fig. 1h-j**; sessions/mice: 27/19 MFB and 13/8 VTA. Quantifications: median ± standard error of the median (s.e._median_)). Successful reaches were 8.3±0.05 mm and 8.4±0.04 mm in length for VTA and MFB respectively (**Fig. 1k**). For comparison, some mice were trained for another 5-7 days to push for water, either before or after DA (15 mice, 5 DA-first). Across quantification metrics, MFB sessions exhibited comparable or higher motivation levels than water sessions, while VTA sessions exhibited comparable or lower motivation levels than MFB and/or water (**Fig. 1h-k**, **Extended Data Fig. 1a**). Both attempted and successful reach rates increased across training, confirming that dopamine reinforcement drives operant learning rather than an acute motor response (successes/min: 5.5±1.2 first day and 8.4±1.5 last day, **Extended Data Fig. 1b-e**). Thus, the push for self-stimulation tasks allowed us to investigate whether and how GrCs and CFs encode DA rewards.

### Imaging cerebellar granule cell and climbing fiber activity during push-for-dopamine

To investigate DA prediction signals in the cerebellum, we focused on its two major input pathways—GrCs and CFs—which constrain downstream cerebellar computation. Recording many GrCs in a local microcircuit requires^27^ transgenic two-photon Ca^2+^ imaging. We also relied on two-photon Ca^2+^ imaging to densely sample CF activity via Purkinje cell dendrites: because there is a well-established 1:1 relationship between climbing fiber action potential bursts and Purkinje cell complex spikes^28^, which in turn exhibit near-perfect concordance (>90%) with dendritic Ca^2+^ transients^29^, the dendritic signals serve as a reliable proxy for climbing fiber input. For simplicity and clarity, we hereafter refer to these Ca^2+^ transients as CF responses. Our GrC imaging (20 mice) employed 3 transgenes (**Fig. 1l**; **Video S3**; Math1-Cre×ztTA×GCaMP6f). For CFs (11 mice), we either used (1) viral R-CaMP2 in Purkinje dendrites of GrC GCaMP transgenics (**Fig. 1m**; 5 mice; AAV-L7-6-R-CaMP2); or (2) mice with transgenic GCaMP in Purkinje cells (**Fig. 1p**, **Video S4**, 6 mice; PCP2-Cre×GCaMP8m).

To combine imaging with push-for-VTA optogenetic activation required several additional workarounds.

Optogenetically activating VTA requires driving opsin expression selectively in DA neurons. Since we already used cre to restrict cerebellar GCaMP expression, an orthogonal genetic driver was needed to restrict opsin to DA cells. We selected DAT-FLP^30^ to virally transduce VTA DA cells with FLP-dependent opsin (AAV-fDIO- ChRimson). Thus, we injected virus and implanted fibers in the VTA of crosses of DAT-FLP with either GrC GCaMP mice (**Fig. 1n**, **Extended Data Fig. 1f**, DAT-FLP×Math1-Cre×ztTA×GCaMP6f), or CF GCaMP mice (**Fig. 1q**, DAT-FLP×PCP2-Cre×GCaMP8m). Additionally, one-photon optogenetics spectrally interfered with the two-photon imaging red channel, which we mitigated (1) via optics (activating ChRimson at 640 nm plus extra emission filters), and (2) by temporally interpolating our R-CaMP2 imaging data around the short 640 nm pulses (**Extended Data Fig. 1h, Methods**). Alternatively, to image during push-for-MFB electrical activation, we simply implanted microwire cannulae targeting the MFB in either GrC- (**Fig. 1o**, **Extended Data Fig. 1g**) or CF-imaging transgenic animals (**Fig. 1r**). We sorted GrCs using nonnegative matrix factorization (cNMF^31^) and computed z-scored fluorescence (**Fig. 1s**, **Extended Data Fig. 1i**). We sorted Purkinje dendrites using independent components analysis (PCA/ICA^32^) (**Fig. 1t**, **Extended Data Fig. 1j,k**), detected spikes via deconvolution and thresholding, and computed spike rates over 150 ms windows. We imaged 100±6 GrCs (cells with detected activity, out of an estimated total visible population of 300-400; all subsequent GrC percentages are relative to the active population), and 35±8 or 127±15 Purkinje dendrites per session with R-CaMP2 or GCaMP respectively (61, 15, and 10 sessions, respectively, **Extended Data Fig. 1l-n**).

In prior studies, cerebellar water reward signals spanned several regions, including Lobule VI^11,33^. Across species and spatial scales, Lobule VI also encodes and contributes to a variety of affective functions^34,35^, as do its target output regions in the cerebellar nuclei^36^. To thoroughly characterize DA reward signals in a consistent cerebellar region, we therefore imaged the right vermis of lobule VI (ipsilateral to the reaching forepaw, contralateral to the self-stimulation sites). We performed imaging studies after 5-7 training sessions.

### Many granule cells sustain anticipatory activity that terminates at DA reward delivery

To examine reward-predictive signals in GrCs, we computed the trial-averaged activity aligned to the end of the reaching movement for each GrC and sorted GrCs by activity level before reward (**Fig. 2a,b**; 20 mice, 14 MFB and 6 VTA). Many GrCs became more active during the delay between the push and the DA reward. Specifically, activity of 18% of GrCs anticipated DA reward, classified by fluorescence significantly elevated just prior to reward and higher than the periods before or after (**Fig. 2c,d**; 100 ms before reward, versus both [400, 700] ms after reward and 300 ms before movement). Inversely, we identified an additional 15% of GrCs that decreased their activity below baseline during the delay (**Fig. 2e,f**). Overall, many GrCs progressively increased or suppressed activity in anticipation of DA reward—despite that no physical action after the initial pushing movement was required to obtain or consume the DA reward.

**Figure 2.**
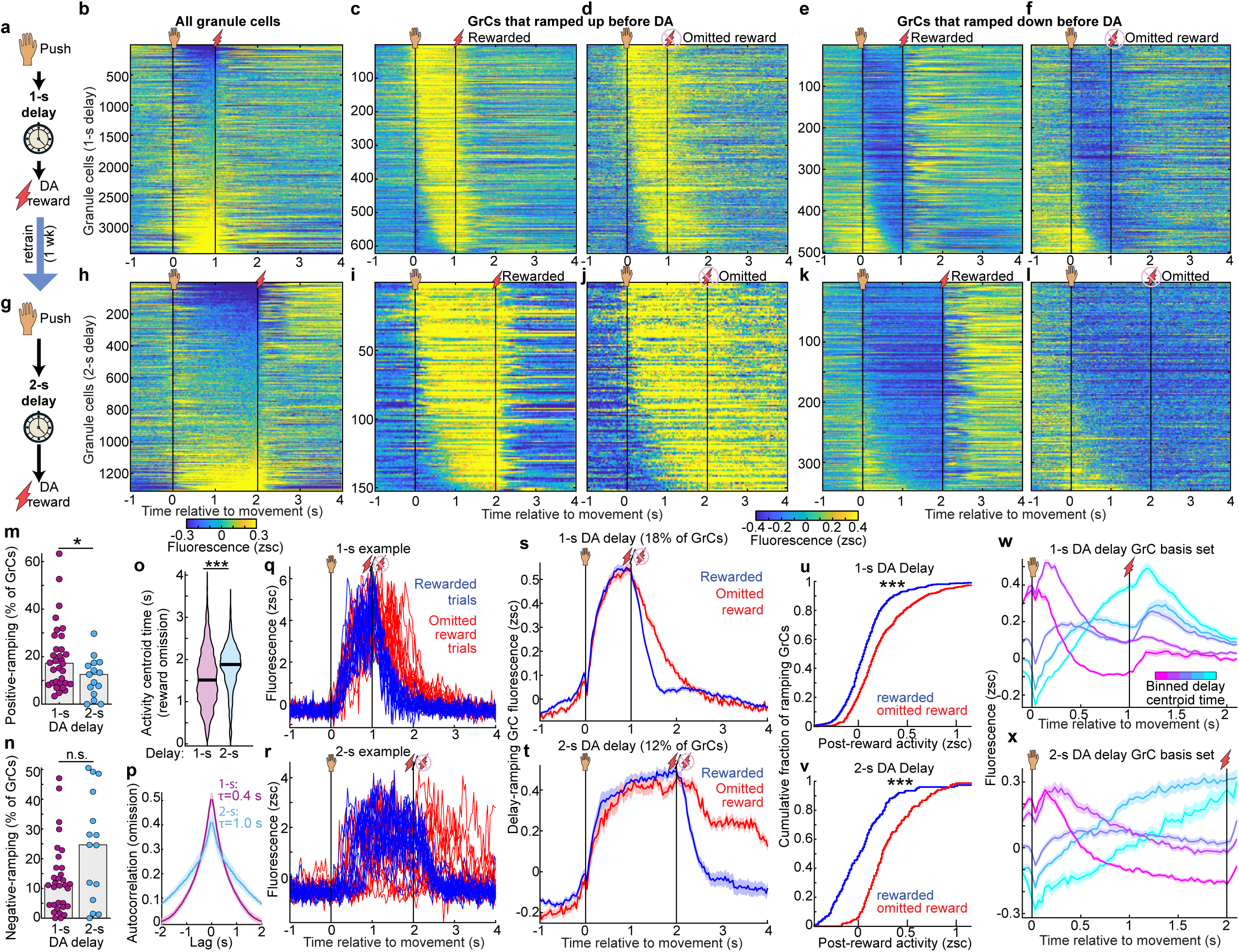
Granule cell activity anticipates dopamine reward and temporally scales to match its expected timing. **a-f,** Push for 1-s-delayed DA task. **a,** Illustration. **b,** Trial-averaged activity of individual GrCs (3,444 cells, 20 mice; 14 MFB/6 VTA). Vertical lines denote movement endpoint and reward delivery. **c-f,** Heatmaps of GrC subsets with positive (**c,d**) or negative (**e,f**) activity ramps before DA reward. (GrC percentages/counts: 18%/621 positive, 15%/502 negative. Positive criteria: activity from [0.9, 1] s significantly elevated and higher than before ([-0.3, 0] s) and after ([1.4, 1.7] s); negative criteria were inverted). **g–l,** Some mice were retrained with 2-s delays (n=9). Panels mirror **a–f** (1,310 GrCs: 12%/152 positive, 26%/347 negative). **m,n,** Per-session percentage of GrCs classified as positive (**m**, p=0.047) and negative delay-ramping (**n**, p=0.06; 32 1-s and 15 2-s delay sessions). **o,** Activity centroid times during omitted reward trials for all GrCs (p<10^-6^). Centroids were timepoints with activity >0.1 zsc between [0, 4] s; criteria were inverted (<-0.1 zsc) for suppressed cells. **p,** Autocorrelation functions averaged across GrCs and sessions on omitted reward trials. Time constants were longer in 2-s delays (*τ*=1.0±0.12 s vs 0.41±0.05 s, median±s.e.m. across sessions; p=0.02; **Extended Data Fig. 2a,b**). **q,r,** 15 rewarded and omitted reward trials from exemplar ramping GrCs in 1-s (**q**) and 2-s delays (**r**). **s,t,** Average activity of all positive-ramping GrCs in 1- (**s**) or 2-s tasks (**t**; 621 GrCs/20 mice and 152 GrCs/9 mice, respectively). **u,v,** Cumulative histograms of post-reward vs post-omission activity across ramping GrCs in 1-s (**u**) and 2-s (**v**) delays (p<10^-6^, mean activity [0.4,0.7] s after reward/omission). **w,x,** Average activity of all GrCs in 1-s (**w**) or 2-s (**x**) delay sessions, binned by their delay activity centroid time (bin edges: 1-s: [-0.1 0.2 0.4 0.55 0.65 0.98] s; 2-s: [-0.1 0.4 0.8 1.2 1.6 1.95]; cell counts per bin: 1-s: [436 907 845 505 572]; 2-s: [265 294 506 144 32]). All comparisons two-sided Mann-Whitney U-test. **m-o,** Centers show medians. **m,n** dots show sessions. Shaded regions show s.e.m. across GrCs.

### Dopamine-anticipating granule cell activity temporally scales to match expected reward timing

We hypothesized that GrC activity ramps “linked” the forelimb movement to the future expected reward. This predicted that longer intervals before DA reward would require longer GrC ramps. To test this possibility, for a subset of animals imaged on 1-s delays, we retrained for another 5-7 days using a 2-s delay (**Fig. 2g**, **Video S5**, 9 mice, 7 MFB and 2 VTA). Visually, most GrC activity profiles appeared temporally “scaled” to the longer delay duration (**Fig. 2h**). Via the same criteria above, we identified 12% of GrCs that ramped activity upwards across the 2-s delay, again with varied kinetics (**Fig. 2i,j**), and 26% of GrCs that ramped activity downwards (**Fig. 2k,l**). Across individual 1-s and 2-s sessions, positive-ramping GrCs comprised 17±3% and 13±3%, while negative- ramping GrCs comprised 11±2% and 25±8% of cells, respectively (**Fig. 2m,n**, 32 1-s and 15 2-s delay sessions).

We next quantified temporal scaling across the entire GrC population. Because delivering rewards at different times will directly drive different activity patterns, we restricted these analyses to omitted-reward trials, where animals experienced no external events after forelimb movement. First, we quantified each GrC by the centroid time of its elevated activity between [0, 4] s on omitted reward trials (**Fig. 2o**). GrC activity was centered substantially later in the trial in 2-s experts than in 1-s experts (p<10^-6^). Alternatively, we considered the temporal autocorrelation function as a metric of the characteristic timescale of GrC activity (**Fig. 2p**). We found that autocorrelation time constants were significantly longer on average in 2-s experts than in 1-s experts (**Extended Data Fig. 2a**, *τ*=1.0±0.12 s vs 0.41±0.05 s, p=0.02). Crucially, this effect persisted even for autocorrelation time constants computed from single trials (**Extended Data Fig. 2b**), confirming that individual GrC transients lengthened rather than only occurring at more variable times across the longer delay.

Suspecting that GrCs ramped for as long as animals continued to await DA reward, we more closely compared rewarded and omitted-reward trials. For both 1-s and 2-s delays; for both positive and negative- ramping GrCs; for individual neurons on single trials (**Fig. 2q,r**, **Extended Data Fig. 2c,d**); and in trial-averages across the population (**Fig. 2s-v**, **Extended Data Fig. 2e**), activity ramps terminated more abruptly following reward delivery, but persisted longer after reward-omission, consistent with active maintenance of an expectant internal state. (By contrast, purely reward-responsive GrCs formed a largely distinct population; **Extended Data Fig. 2f,g**). By binning the entire GrC population based on delay activity timing, we found that ascending and descending ramps of varying kinetics all roughly doubled in duration in 2-s vs 1-s delays (**Fig. 2w,x**). Thus, GrCs ramped up and down with heterogeneous profiles that temporally scaled to link the action to the expected time of DA reward delivery.

### GrC populations represent expectation of DA and water rewards similarly

Most prior studies of reward signaling in cerebellar circuits relied on water reward in thirsty animals—a reward that is both homeostatic and ethologically relevant, but complicated by extensive orofacial drinking movements. To determine the generality of GrC reward representations in individual animals, a subset of mice imaged in push-for-DA were also imaged in push-for-water (**Fig. 3a**, **Video S6**, 10 mice, 6 trained in water first, 4 in DA first; 6 MFB and 4 VTA). We analyzed this dual-context dataset in two sequential stages: first evaluating whether the *population-level* prevalence and magnitude of anticipatory activity were conserved across reward types (**Fig. 3**), and subsequently tracking whether *individual* GrCs generalized their representations across these distinct contexts (**Fig. 4**). In both water and DA reward tasks, these mice exhibited substantial delay-period GrC activation (**Fig. 3b,c**). GrCs with elevated activity prior to reward comprised similar proportions of the population in DA (18%) and water (14%) reward contexts, both with a variety of temporal kinetics (**Fig. 3d–g**). Similarly, negative ramping cells comprised 14% and 17% of GrCs in DA and water, respectively (**Extended Data Fig. 3a,b**). Across individual imaging sessions, positive delay-ramping cells comprised 18±5% (DA) and 10±2% (water) of GrCs (**Fig. 3h**), while negative ramping cells comprised 11±4% (DA) and 12±3% (water) of GrCs (**Fig. 3i**). Activity levels of ramping GrCs just prior to reward were slightly higher for DA than water (**Fig. 3j**). By comparing mice trained first with water reward to those trained first with DA, we confirmed that DA anticipation signals did not require prior experience with water reward (**Extended Data Fig. 3c-f**). For both water and DA rewards, GrC ramps terminated more rapidly following reward than reward omission (**Fig. 3k-n**), suggesting that ramps persist while animals await reward, regardless of its type. Thus, GrC populations exhibit comparably strong and widespread predictive encoding of DA rewards and homeostatic water rewards.

**Figure 3.**
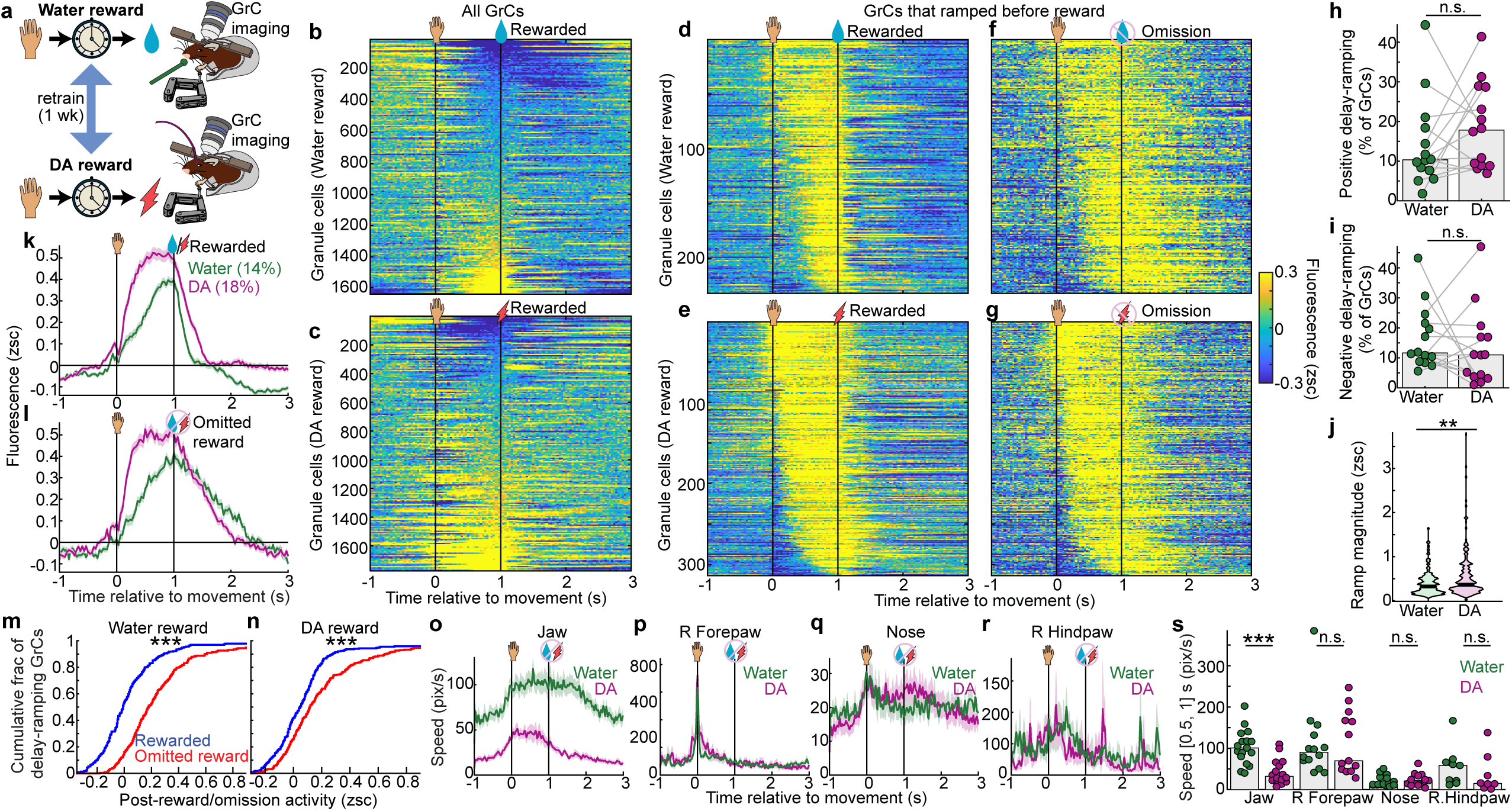
GrC populations represent expectation of DA and water rewards similarly. **a**, Experimental timeline. GrCs were imaged across sequentially trained push-for-water and push-for-DA tasks (5-7 training sessions each; 10 mice, 6 water-first and 4 DA-first; 6 MFB and 4 VTA). **b,c,** Trial-averaged activity of all GrCs in water (**b**) and DA (**c**) tasks. Each raster sorted independently by pre-reward activity ([0.9, 1] s; GrCs: 1,660 water and 1,754 DA). **d–g**, Heatmaps of GrCs exhibiting positive pre-reward ramping (defined as in Fig. 2), averaged across either rewarded (**d,e**) or omitted reward (**f,g**) trials in water (**d,f**; 231 GrCs) or DA (**e,g**; 313 GrCs) tasks. Cells are sorted by temporal centroids of elevated activity (>0.1 zsc) during the delay. **h,i,** Per-session prevalence of positive (**h**, p=0.3) and negative (**i**, p=0.4) delay-ramping GrCs (14 DA and water sessions from 10 mice). **j,** Average pre-reward activity ramp magnitude for all ramping GrCs ([-0.1, 0] s from reward; p=0.002, GrCs: 231 water and 313 DA). **k,l,** Population mean activity traces of positive-ramping GrCs for rewarded (**k**) and omitted reward (**l**) trials (shaded regions show s.e.m. across GrCs). **m,n,** Cumulative histograms of post-reward vs post-omission activity across ramping GrCs in water (**m**) and DA (**n**) tasks. ([1.4, 1.7] s, both p<10^-6^). **o-s**, Body movement. **o-r**, Average body speed of the jaw (**o**), right forepaw (**p**), nose (**q**), and right hindpaw (**r**) on omitted reward trials (traces show mean and shaded regions show s.e.m. across session-averages. **s,** Mean pre-reward speed ([0.5 1] s). Bonferroni-corrected p-values: jaw = 0.0007; others: 1. Sample sizes (mice/water sessions/DA sessions): jaw (13/17/16), right forepaw (11/14/13), nose (12/16/15), right hindpaw (6/9/8). Dots show sessions. Bars show medians. **h,i**, two-sided Wilcoxon sign-rank test (14 matched session pairs). **j,m,n**,**s,** two-sided Mann-Whitney U test.

**Figure 4.**
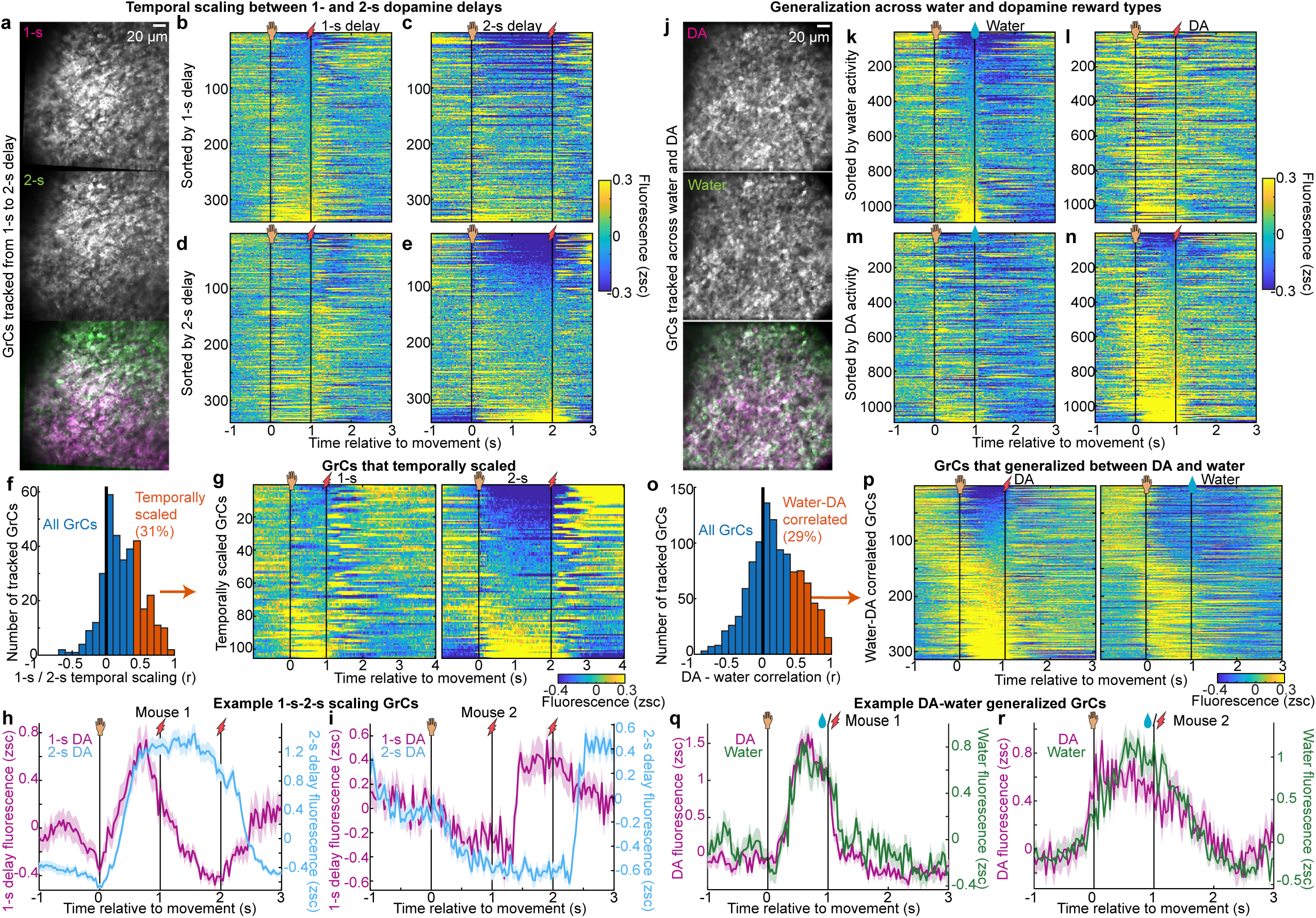
Many individual GrCs generalize across action-dopamine delay durations or reward types. **a-i,** GrCs tracked across 1-s and 2-s DA delay sessions (5-7 days apart). **a,** Example 1-s-2-s-matched imaging field. **b-e,** Trial- averaged activity on rewarded trials for all tracked GrCs, sorted by 1-s (**b,c**) or 2-s (**d,e**) delay activity (340 cells from 5 fields in 4 mice). **f,** Temporal scaling index distribution. Index was the Pearson correlation (*r*) between a neuron’s 1-s profile and its temporally compressed 2-s profile ([-1,3] s; only delay period compressed). 31% of neurons were classified as temporally scaled (*r* >0.4, Benjamini-Hochberg 1% false discovery rate [FDR]). **g,** Trial-averaged activity for the 104 temporally scaled GrCs. **h,i,** Example temporally scaled GrCs from two mice (traces show mean±SEM across trials). **j-r**, GrCs tracked across water and DA reward sessions (5-7 days apart). **j,** Example imaging field matched across water and DA sessions. **k-n,** Trial-averaged activity for all tracked GrCs (1,098 GrCs from 11 fields in 8 mice), sorted by water (**k,l**) or DA (**m,n**) activity. **o,** Water-DA correlation distribution. Index is the Pearson correlation (*r*) between a neuron’s water and DA activity profiles ([-1,1] s excluding a window around the forelimb movement, [-0.2, 0.1] s). 29% of neurons generalized across reward types (*r* >0.4, 1% FDR). **p,** Trial-averaged activity for the 314 generalized GrCs. **q,r,** Example generalized GrCs from two mice**. h,i,q,r,** Traces show means and shaded regions s.e.m. across trials.

### DA reward-anticipating GrC dynamics are incompatible with motor kinematics

In contrast to water reward which elicits licking both before and during reward consumption, no physical action is needed to consume the DA reward—yet animals might still compensate via new uninstructed anticipatory movements. To test this, we tracked body movement using DeepLabCut^37^ and found that, compared to water, DA sessions elicited 68% less anticipatory jaw movement (**Fig. 3o,s** 101±11 vs 32±9 pix/s, p=0.0007 Bonferroni-corrected), without adding other anticipatory body movements (**Fig. 3p-s**, all p=1). Accordingly, across matched imaging fields, the change in anticipatory jaw speed between water and DA reward was unpredictive of the change in prevalence of anticipatory GrCs (p=0.63, **Extended Data Fig. 3g**). To dissociate other residual movements from DA reward-anticipating GrC signals, we leveraged the abrupt cognitive change from reward expectation to receipt as a key dissociation. Comparing rewarded and omission trials revealed a double dissociation: after reward delivery, physical movement spiked (consistent with known stimulation-induced movements^38^) but anticipatory GrC activity rapidly quenched; conversely, reward omission elicited no increase in movement, yet GrC activity persisted (**Extended Data Fig. 4a-c**). A similar double dissociation emerged from comparing the 1-s and 2-s delay conditions during the 1.5–2 s window: having just received reward, the 1-s cohort exhibited elevated movement but quenched neural activity, whereas the 2-s cohort exhibited minimal movement but peak neural anticipation due to the ongoing delay (**Extended Data Fig. 4d-f**). To formalize this decoupling in individual neurons, we used variance partitioning. We regressed each anticipatory GrC’s activity (concatenating the rewarded and omitted-reward trial-averages) simultaneously against both a model of anticipatory ramping and the body kinematics (two principal components, **Methods**). Because purely kinematic models fail to capture the cognitive difference between reward expectation and receipt, GrCs most strongly quenched by reward had lower proportions of variance independently explained by movement (**Extended Data Fig. 4g-k**). Importantly, anticipatory representations—and their dissociation from post-reward kinematics—were structurally conserved across both VTA and MFB cohorts, with MFB reinforcement eliciting larger, more prevalent neural ramping signals and also driving stronger motivation (**Extended Data Fig. 5**). Finally, to demonstrate that instrumental action was not required to initiate reward-predicting GrC ramping, we imaged another cohort trained in a passive DA anticipation paradigm (7 MFB mice). Even in this passive context, a purely auditory cue predicting 1-s-delayed DA reward elicited anticipatory ramping in 24% of GrCs with strength and timing comparable to the operant condition (**Extended Data Fig. 6**). Collectively, these data demonstrate that GrC anticipatory ramps reflect the genuine internal expectation of reward, featuring dynamics that are qualitatively incompatible with residual body movements and independent of instrumental motor requirements.

### Individual GrCs temporally scale with DA delay duration and generalize across water and DA reward

Having demonstrated a common GrC population representation across both delay durations and reward types, we questioned whether individual GrCs also generalized: either by temporally scaling across 1- and 2-s delays; or by cross-activating for water and DA. In some 2-s-delay recordings, we were able to register individual GrCs to recordings 5-7 days prior with a 1-s delay (**Fig. 4a**). We then restricted analysis to GrCs matched across 1- and 2-s-delay recordings, and again generated trial-averaged raster plots (**Fig. 4b-e**, **Extended Data Fig. 7a** 340 GrCs from 4 mice, 2 MFB and 2 VTA). Visually, only a fraction of cells had activity that seemed similar across delay durations. To identify individual GrCs with activity that temporally “stretched” to the delay duration, we compared each neuron’s 1-s-delay activity to a “temporally scaled” version of its 2-s-delay activity, by compressing the activity during the 2-s delay into a 1-s span, and then computing the Pearson correlation coefficient on the resulting activity traces (**Fig. 4f**). To conservatively classify generalizing GrCs in both the 1- s/2-s and water/DA datasets, we applied a minimum effect size threshold (*r* > 0.4) derived from a False Discovery Rate correction (*q* < 0.01, see **Methods**). These criteria scored 31% (104 of 340) of all registered GrCs as temporally scaled (**Fig. 4g**; shuffle-controls: 5% **Extended Data Fig. 7b**). Some of these GrCs, for example, exhibited elevated activity during the 1-s delay that was stretched to match the longer delay in the 2-s-delay recording (**Fig. 4g,h**, **Extended Data Fig. 7c**). Other GrCs exhibited reduced activity during the 1-s delay that was extended into longer suppression in the 2-s-delay recording (**Fig. 4g,i**, **Extended Data Fig. 7d**). In the remaining non-scaling population, a trial-by-trial reliability analysis revealed that some GrCs encoded only a single delay context, while others reliably encoded both contexts but with distinct, non-scaling activity profiles (**Extended Data Fig. 7i**). Thus, many GrC activity profiles temporally scaled to the interval between action and DA reward time, without correspondingly extended body movements (**Extended Data Fig. 4**), suggesting that such cells represent reward temporal expectations.

We next examined whether individual GrCs played similar roles across both DA and water reward types. By registering neuronal identity across DA and water sessions (**Fig. 4j**), we matched 1,098 GrCs from 8 mice across both contexts (**Extended Data Fig. 7e**, 4 MFB and 4 VTA). When sorting these GrCs based on their activity level just prior to water (**Fig. 4k,l**) or DA (**Fig. 4m,n**), we observed only a subset of GrCs with similar activity profiles in both contexts. To identify neurons with cross-reward generalized activity, we computed for each GrC the correlation from [-1,1] s between its trial-averaged activity profile in water and that in DA (**Fig. 4o**, excluding 300 ms around the common reaching movement [-0.2, 0.1] s). The cutoffs applied for temporal scaling also classified 29% (314 of 1098) of all water-DA registered GrCs as similar (shuffle controls: 15%, **Extended Data Fig. 7f**). Thus, a subset of GrCs also generalized across reward types (**Fig. 4p-r**, **Extended Data Fig. 7f- h**), despite the far greater anticipatory movements in the water context (**Fig. 3o**), suggesting such cells may encode more abstract features of reward-driven behavior.

### Many climbing fibers respond at short latency to dopamine reward delivery

Synaptic integration of GrC inputs in Purkinje cells is dictated partly by their relationship to near-coincident CF inputs^39,40^. To gain insight into the relationship between GrC and CF reward dynamics, we imaged CF activity (reported by Purkinje dendritic Ca^2+^ spiking) during push-for-DA (**Fig. 5a**, 11 mice, 6 MFB and 5 VTA). Sorting CFs by their activity just after self-stimulation delivery (spike rate [0.1, 0.2] s), we noted numerous neurons with short-latency responses to DA reward (**Fig. 5b**, 1,048 CFs). We classified CFs as DA reward-responsive if their mean spike rate [0.1, 0.2] s was >1.3 Hz (**Fig. 5c**, 66% of CFs). We noted that such responses were absent when reward was omitted. Disaggregating by MFB versus VTA reinforcement confirmed that DA reward-responsive CFs exhibited conserved, robust response magnitudes, but trended toward greater prevalence during MFB sessions (**Extended Data Fig. 8a-f**), paralleling their higher behavioral motivation levels (**Fig. 1**). To determine whether this DA reward-evoked CF activity acts as an initial instructive signal rather than a learned prediction, we examined responses in naive mice on Day 1 of training. We found that widespread, robust CF responses to DA reward are present from the very outset of learning (**Extended Data Fig. 8g,h**), confirming their capacity to serve as instructive signals.

**Figure 5.**
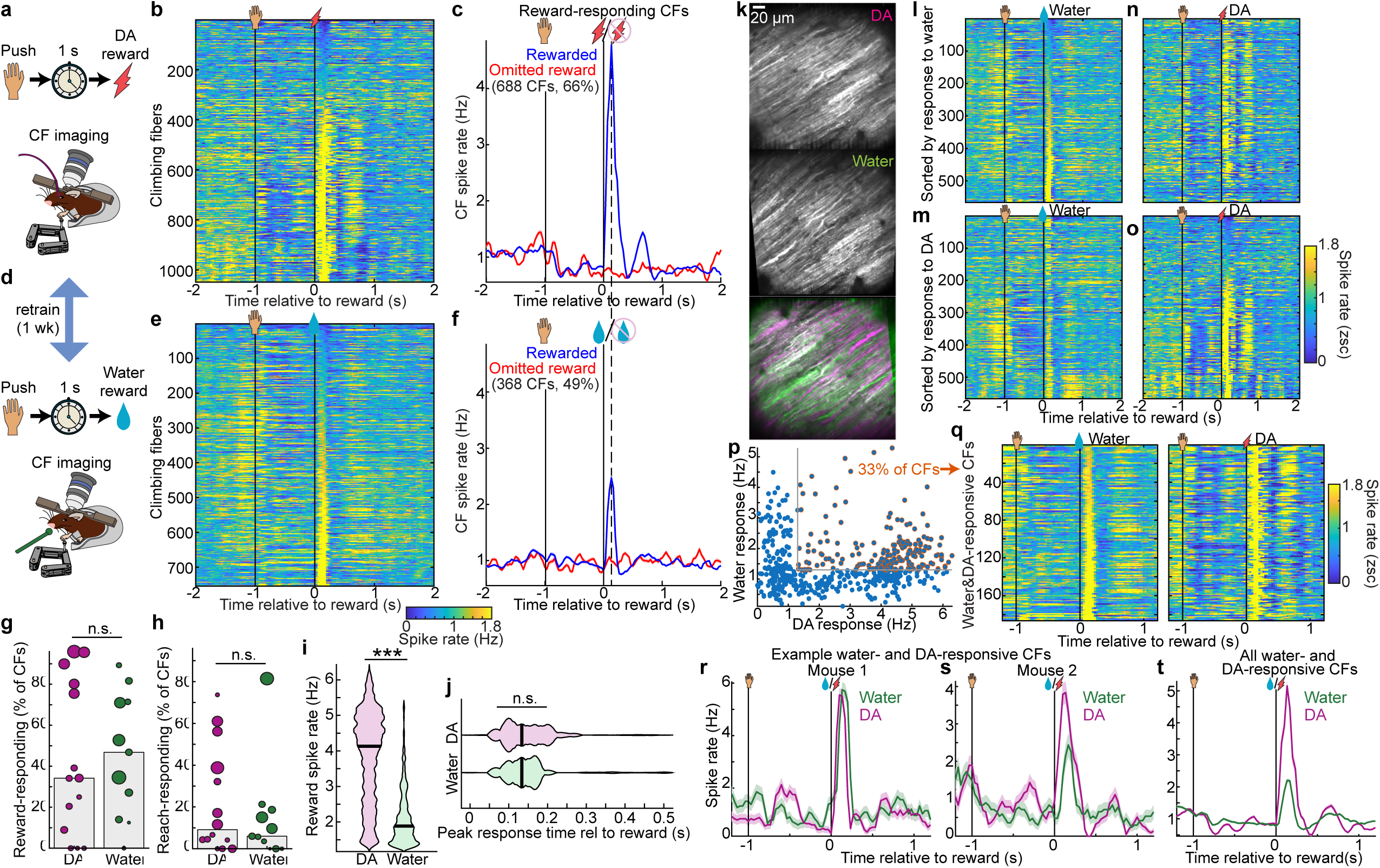
Dopamine rewards activate most CFs, many of which also spike after water rewards. **a-c**, CF imaging during DA and water reward tasks. **a,** DA task schematic. **b,** Trial-averaged activity of individual CFs aligned to DA reward delivery (1,048 CFs from 11 mice). **c,** Population average of the 688 CFs (66%) classified as DA reward-responsive (post-reward spike rate >1.3 Hz, from [0.1, 0.2] s). **d-f**, Parallel data for the water reward task (754 CFs from 8 mice), showing task schematic (**d**), individual CFs (**e**), and the population average for 368 water reward-responsive CFs (49%). **g,h**, Per-session prevalence of reward-responding (**g**, p=0.8) and reach-responding CFs (**h,** p=0.6). Reach responsive was defined as spike rate >1.3 Hz from [-0.1, 0.1] s relative to movement. Area of each session’s dot is proportional to number of CFs. **i,j,** Violin plots comparing the post-reward spike rate ([0.1, 0.2] s p<10^-6^) (**i**) and peak response time (p=0.2) (**j**) for all reward-responding CFs (688 DA and 368 water CFs). **k-t**, CF tracking across water and DA sessions. **k,** Example fields showing CF registration. **l-o,** Trial-averaged activity of tracked CFs (566 CFs from 8 mice) in water and DA tasks, sorted either by their water reward response (**l,n**) or DA reward response (**m,o**). **p,** Scatter plot of each cell’s mean post-reward activity ([0.1, 0.2] s) in DA vs water sessions. 33% of tracked CFs were reward-responsive in both contexts (>1.3 Hz). **q-t,** Activity of the 186 water- and DA-responsive CFs. **q,** Trial-averaged heatmaps sorted by water response. **r,s,** Example water- and DA-responsive CFs from two mice (traces show mean ± s.e.m. across trials). **t,** Population average across all 186 water- and DA-responsive CFs. **g-j,** Centers show medians and comparisons are two-sided Mann-Whitney U-test. **c,f,t,** Traces show means and shaded regions s.e.m across CFs. **r,s**, traces show means and shaded regions s.e.m. across trials.

Having previously observed similar responses during a water reward task, we compared these reward responses for DA to those for water in a subset of mice trained in both tasks (**Fig. 5d**, 8 mice, 4 MFB and 4 VTA; 7 trained on water first). Water reward responses (**Fig. 5e**), though comparable to those we reported previously^11^, were far smaller than those seen for DA rewards. Quantitatively, water reward-responsive CFs by the above cutoff comprised 49% of cells, but their response spike probabilities were far lower than for DA rewards (**Fig. 5f**). On a per-session basis, DA reward-responding and water-responding cells comprised a comparable 34±17% and 47±16% of CFs, respectively (**Fig. 5g**, p=0.8). By comparison, applying the same criteria to classify reach- responding cells identified only 9±10 and 6±5% of cells in DA and water sessions, respectively (**Fig. 5h**). Spike rates in response to DA reward (4.1±0.06 Hz) far exceeded the response to water reward (1.9±0.04 Hz; **Fig. 5i**), but peak response latencies were the same: 134 ms (**Fig. 5j**). Thus, many CFs responded rapidly following DA rewards with far higher probability than typical responses to water rewards.

To determine whether individual CFs responded to both water and DA rewards, we registered Purkinje dendrite identities across tasks (**Fig. 5k**). Visually, we noted limited overlap when sorting CFs based on water reward responses (**Fig. 5l,n**) or DA reward responses (**Fig. 5m,o**). To compare each CF’s DA and water reward responses, we scattered each cell’s mean spike rate from [0.1,0.2] s from the two recordings (**Fig. 5p**). Roughly 1/3^rd^ of CFs cleared the same 1.3 Hz cutoff for both DA and water rewards (**Fig. 5q,r**). However, in contrast to the far above-chance GrC generalization, and likely due both to the high correlations among groups of CFs^41^ and the very high proportion of DA reward-responsive CFs overall, dual reward-responsive CFs were not substantially more common than in shuffle controls (p=0.06, **Extended Data Fig. 8i**). In addition, even CFs responsive to both water and DA rewards responded far more strongly to DA rewards (**Fig. 5s,t**, **Extended Data Fig. 8j,k**). Overall, DA rewards elicited widespread spiking responses in CFs, which also often responded to water reward but more weakly.

### Naïve mice learn to push at moderate rates for delayed CF activation, with predictive GrC ramping

If CF reward signals are instructive cues for decoding reward-predicting GrC activity, they might also drive motivated operant learning. To test this, we trained naïve mice to push solely for delayed CF activation ‘rewards.’ We used transgenic GrC GCaMP mice (Math1-Cre×ztTA×GCaMP6f) injected with the red-shifted opsin ChRmine in the inferior olive to activate posterior cerebellar climbing fibers through an imaging cranial window (**Fig. 6a,b**; 10 CF reinforcement mice; 4 no-opsin light-only control mice). To simulate the observed CF reward response, we delivered one 50-ms pulse of 640 nm light centered over Lobule VI triggered 1 s after successful reach end (**Fig. 6c**). After 5-7 training sessions, CF-reinforced mice executed 5.9±0.4 successful and 6.6±0.3 attempted pushing movements per minute, yielding an overall completion rate of 89±2% (**Fig. 6d–g**; light-only controls after training: 2.3±0.3 successes/min, 3.4±0.6 attempts/min, and 68±11% completion). This indicates a surprising capacity for this pathway to drive behavioral reinforcement, although overall motivation remained lower than with primary dopaminergic or water rewards (**Extended Data Fig. 9**). To confirm that this performance reflected *bona fide* associative learning rather than an acute, non-specific increase in movement, we tracked behavioral metrics across the training period (5–7 sessions). While both attempted and successful reach rates progressively increased for CF-reinforced mice, performance in light-only controls steadily degraded (**Fig. 6h– j**). This demonstrates that delayed CF activation is sufficient to reinforce the *de novo* acquisition of the operant behavior, whereas control animals adaptively extinguish the unrewarded action.

**Figure 6.**
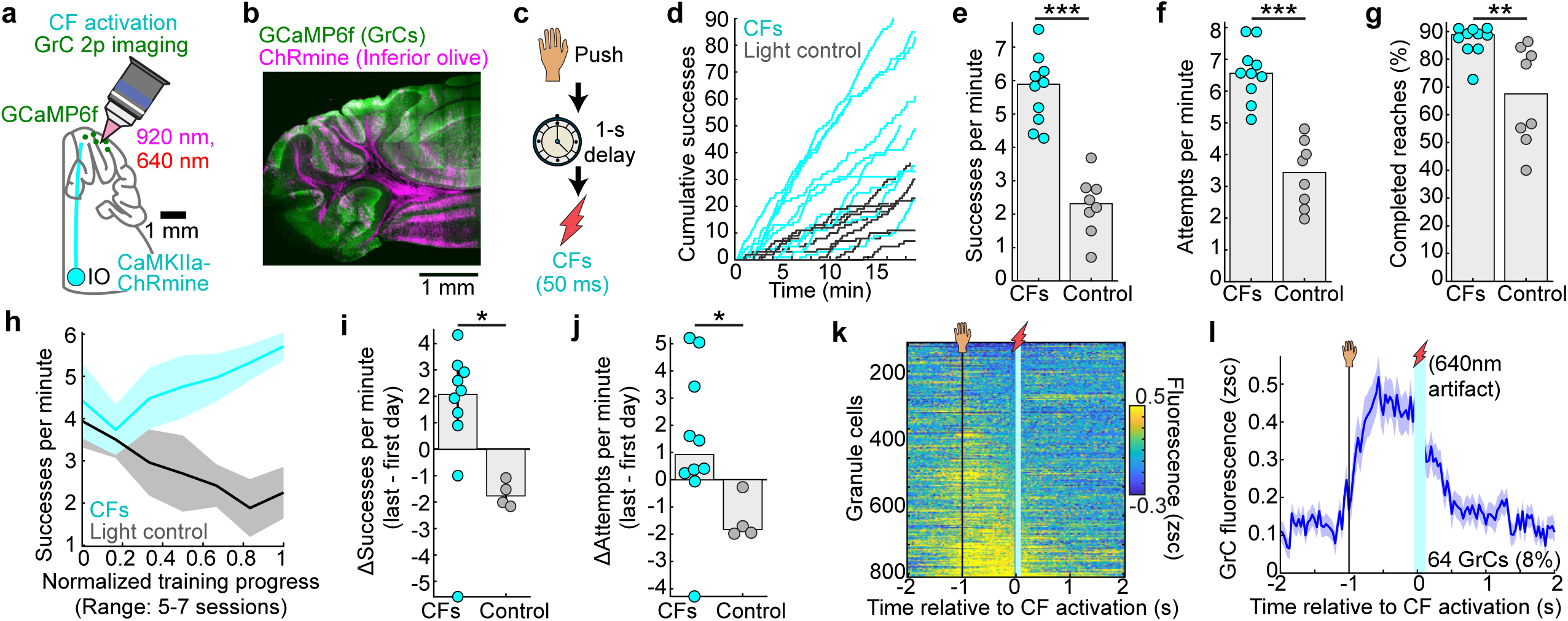
Naïve mice learn to push at moderate rates for delayed CF activation, with predictive GrC ramping. **a-g**, CF activation via ChRmine in the inferior olive (IO) of GrC GCaMP transgenics. **a,** Task schematic. **b,** Histology showing GrC GCaMP and CF ChRmine expression. **c,** Paradigm (training proceeded for 5-7 sessions). **d-g,** Behavioral metrics comparing CF reinforcement to light-only controls. **d,** Cumulative successful reaches over time for each session. **e-g,** Successes per minute (**e**), attempts per minute (**f**), and completion percentage (**g**) (**e-g** p values: 5×10⁻⁵; 5×10⁻⁵; 0.003; sessions/mice: 10/10 opsin and 8/4 light-only control). **h-j,** Behavior metrics over training. Successes per minute (**h,i**) and attempts per minute (**j**) increased more for CF reinforcement than light controls (both p=0.02). Lines and shaded regions in **h** show mean and s.e.m. across mice. **k,l,** Neural activity during CF stimulation. **k,** Rasters of all GrCs aligned to CF stimulation (812 GrCs from 10 sessions/mice). **l,** Average activity of GrCs that ramped prior to expected CF activation (8%, 64 delay-ramping GrCs, classified as in Fig. 2; traces show mean and shaded regions s.e.m. across GrCs). **e–g,** Bars show medians and dots show sessions. **i,j,** Bars show medians and dots show mice. Comparisons are two-sided Mann–Whitney U-tests.

We next examined GrC responses during push-for-CF activation. Because the 640 nm light partly contaminated the green imaging channel, we discarded imaging data during the 50-ms pulse of CF activation light. We found that a small but notable group of GrCs appeared to show elevated activity during the delay period prior to CF stimulation (**Fig. 6k**). Examining this group of neurons in more detail via the criteria from **Fig. 2**, we found delay ramping activity similar to that seen in push-for-reward sessions, but with lower prevalence (**Fig. 6l**, **Extended Data Fig. 9f**). Thus, reward-evoked CF activity can reinforce associative operant behavior, while similarly eliciting GrC activity ramps that link the action and future stimulation outcome.

### Perturbing delay-period GrC activity throughout training disrupts push-for-DA learning

We next evaluated the causal role of action-DA interval GrC activity for associative learning. We used Math1- Cre × LSL-stGtACR1 mice to provide randomly timed optogenetic GrC silencing selectively during the delay period throughout push-for-MFB learning (**Fig. 7a,b**; **Extended Data Fig. 10a**). While this strategy cannot exclude a role for GrC activity in other task epochs, delay-period inhibition specifically targets the window where the anticipatory signal natively emerges (**Fig. 2**). Furthermore, restricting inhibition to the period immediately following reach completion minimizes direct motor effects on the subsequent reach (>4 s later). One group of Math1-Cre×stGtACR1 animals received a single pulse of light centered on Lobule VI during the delay period before MFB reward on every trial for 5-7 days (11 mice). To ensure the optogenetic perturbation could not serve as a fixed temporal cue, we randomized both the laser onset during the delay period and its duration on a trial-by-trial basis (median±m.a.d.: onset, 300±180 ms after pushing >6 mm; duration, 370±84 ms; **Fig. 7c,d**). A second group of littermate controls received normal (i.e., no-laser) training (8 mice). To establish a statistically powered normative baseline, we pooled data from the 8 no-laser Math1-cre×stGtACR1 mice with the 27 MFB imaging mice. Both cohorts underwent identical surgical and training procedures and exhibited indistinguishable behavioral performance (**Extended Data Fig. 10b–d**). Compared to this normative baseline, mice trained with GrC delay-period inhibition executed significantly fewer successes and attempts per minute, yielding lower overall completion rates (**Fig. 7f–h**). To confirm that these deficits reflected impaired task acquisition rather than solely acute motor interference, we evaluated the cohorts using two independent approaches. First, longitudinal tracking revealed that while baseline mice progressively improved their performance across training days, GrC- inhibited mice failed to show similar associative gains (**Extended Data Fig. 10e–g**). Second, these behavioral impairments only partially recovered during the first laser-off "washout" session, confirming that the learning deficits outlasted the acute optogenetic perturbation (**Extended Data Fig. 10h–j**). Thus, intact GrC delay-period activity is necessary to acquire and sustain the motivated vigor needed for operant learning.

**Figure 7.**
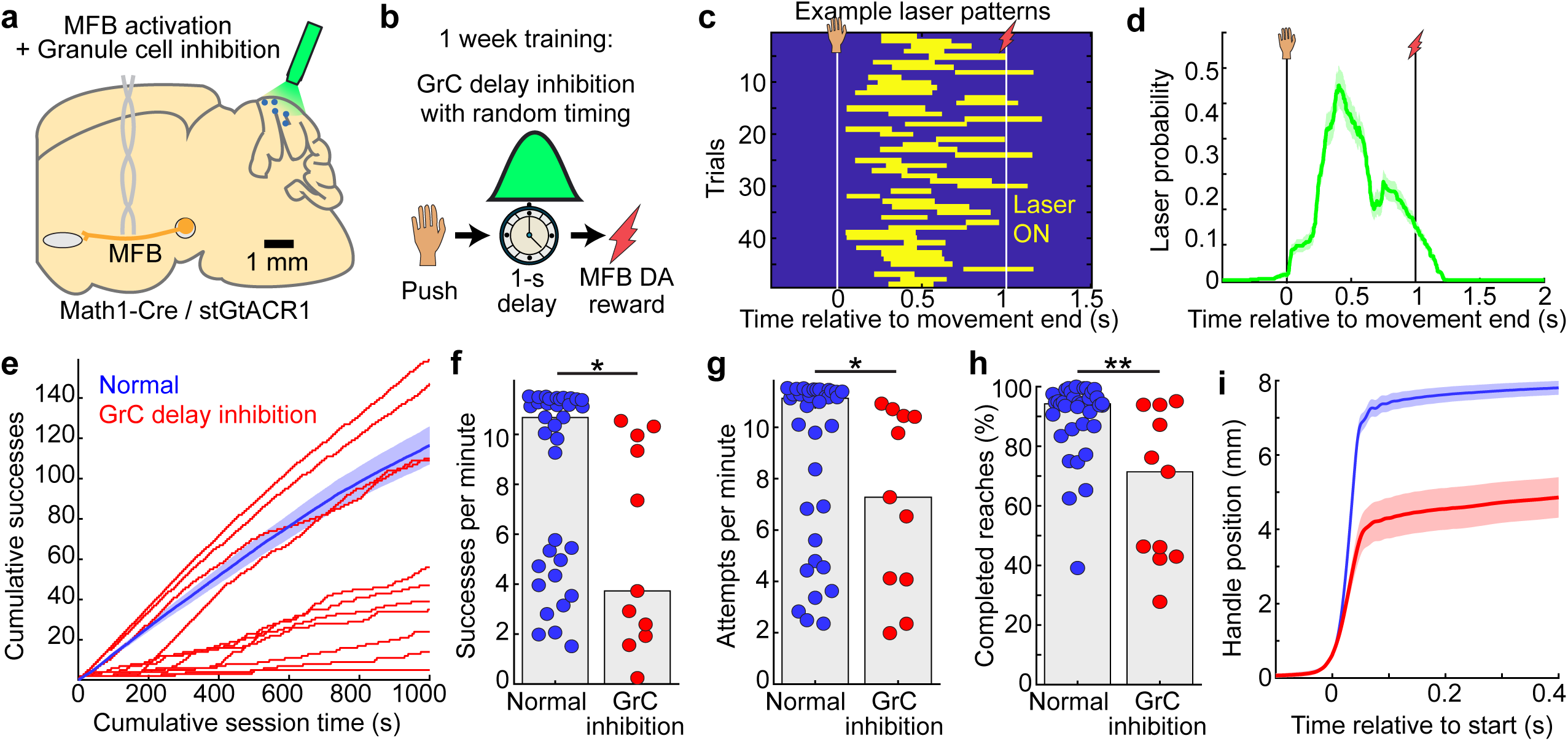
Perturbing delay-period GrC activity throughout training disrupts push-for-DA learning. **a,b,** Optogenetic GrC inhibition during MFB reward training (5-7 sessions). **a,** Strategy for stGtACR1-mediated GrC inhibition. **b,** Task paradigm. GrCs were inhibited during the 1-s delay period using randomized laser timing to prevent the light from acting as a learnable cue. **c,d,** Distribution of laser activation times for an exemplar session (**c**) and averaged across all inhibition sessions (**d**, 11 sessions/mice). **e,** Cumulative successful reaches over time for each session. The “Normal” control group (35 sessions) underwent the same surgery as the inhibition cohort and includes 27 MFB sessions from 19 imaging mice and 8 MFB sessions from 8 genotype-matched stGtACR1 controls (blue trace shows mean and shaded region s.e.m. across sessions; subgroups were behaviorally equivalent, **Extended Data Fig. 10b-d**). **f-h,** Per-session behavioral metrics. **f,** Successes (p=0.02) and **g,** attempts (p=0.04) per minute. **h,** Completion percentage (p=0.006). 35 normal and 11 GrC inhibition sessions. p-values are Bonferroni-corrected two-sided Mann-Whitney U-test, dots show sessions, and bars show medians (comparison to laser-off test days in **Extended Data Fig. 10h-j**). **i,** Handle position aligned to movement start (mean across session-averages). **d,i,** Traces show mean and shaded region s.e.m. across sessions.

## DISCUSSION

By imaging cerebellar GrCs and CFs in mice pushing for delayed dopaminergic self-stimulation, we found that these input streams conveyed robust predictive and instructive representations of expected dopamine rewards. Rather than merely coordinating the kinematics of consumption, GrC populations tracked the internal expectation of the abstract reward, while time-locked CF activity signaled its arrival (**Fig. 5**). GrCs generated heterogeneous activity ramps that temporally scaled to meet task demands, successfully linking expected DA reward back to the instrumental action (**Fig. 2**) or passive sensory cue (**Extended Data Fig. 6**). Crucially, this representational strategy was conserved within individual mice sequentially trained across DA and water rewards, despite the far greater anticipatory movements before water (**Fig. 3**, **5**). In fact, many individual GrCs (**Fig. 4**) and CFs (**Fig. 5**) exhibited cross-activation in both water and DA contexts. This suggests that the cerebellar cortex maintains state-like representations of reward expectation that transcend specific sensorimotor constraints. Causally, delayed CF activation on its own (in place of reward) drove moderate associative operant learning in naïve mice (**Fig. 6**)—to our knowledge the first demonstration that cerebellar reward signals suffice to reinforce a novel action (complementing their necessity for water-driven learning^42^). Conversely, delay-period GrC activity was required to learn and execute the task with a normal degree of motivation and vigor (**Fig. 7**). Together these results suggest deeper integration of cerebellar circuits into reward prediction networks than previously understood.

The temporal distribution of ascending and descending anticipatory GrC activity ramps forms a temporal “basis set”^43,44^ (**Fig. 2w,x**), which is a central postulate of cerebellar temporal prediction theory^45^. In this study, we demonstrated that GrC temporal basis sets are present even for abstract events like self-stimulation rewards. The basis set exhibits several computationally important features. First, “temporal scaling” enabled GrCs to span the interval from action to the expected time of reward, precisely when CF feedback signals arrived. By bridging this gap, GrC ramps remain “visible” to CF-dependent plasticity mechanisms at downstream Purkinje cell synapses^46^, effectively solving the local credit assignment problem for delayed rewards. Second, many basis elements (individual GrCs) generalized between anticipation of DA and water rewards, despite their divergent sensorimotor requirements. Similarly, many CFs spiked after both water and DA rewards, reminiscent of previous results in which CFs cross-activate to different events^28^ depending on salience^14^. This cross-contextual stability may permit the cerebellar cortex to compute general timing signals when knowledge of one context applies in another, potentially facilitating coordination in multimodal contexts^47,48^. At the same time, the substantial group of GrCs exhibiting context-specific activity could support environmental disambiguation. Future studies will be required to determine how these cerebellar representations adapt to more complex reinforcement landscapes, such as variations in outcome magnitude, unpredicted reinforcement delivery, or the systematic degradation of temporal predictability. While our data demonstrate robust predictive and instructive DA reward encoding at the cerebellar input layer, the cerebellar computations performed on these inputs downstream, and how they mediate the causal behavioral effects observed here, remain critical open questions.

Replacing reward with CF activation sufficed to drive moderate operant learning (**Fig. 6**), clearly indicating causal cerebellar involvement in reward-driven associative learning. While these data suggest that cerebellar circuits reinforce self-initiated actions in a more limited way than the nigrostriatal system (**Extended Data Fig. 9** CFs vs VTA/MFB), the observed level of CF reinforcement likely represents a lower bound, given technical constraints inherent to the approach: (1) we transduced a minority of CFs; (2) fiberoptics illuminate a small fraction of the total volume (∼1mm^3(49)^, or ∼2%^50^); and (3) bulk stimulation activates functionally heterogeneous CFs. Given the paramount role of the nigrostriatal system in driving motivated behavior^51^, and the extensive projections from the cerebellar nuclei to both the VTA monosynaptically^52^ and the striatum disynaptically^53^, determining whether these effects operate via the dopaminergic system is a critical direction for future research.

Conversely, chronic perturbations of GrC delay activity disrupted push-for-DA learning (**Fig. 7**), in ways qualitatively similar to studies of consummatory rewards^11,42^. Specifically, GrC inhibition degraded multiple components of the operant loop, including success rates and attempt frequencies. Drawing on prior literature, we propose these behavioral outcomes reflect complementary mechanisms rather than mutually exclusive functions: anticipatory GrC activity likely helps reinforce the preceding action^16^, shortens the reaction time for subsequent attempts^54^, and sustains the motivated state required for task engagement. Still, resolving the role of different GrC basis set elements and their temporal patterning will require further technical advancements in two-photon optogenetics. Further work will also be needed to delineate the specific role of the cerebellum in reward-driven learning compared to other brain networks like the nigrostriatal system—essential to building a holistic brain-wide model of reward-based learning.

Furthermore, several findings in this study bear on the relationship between motivation and cerebellar dynamics. VTA DA reward drove weaker anticipatory GrC signals than MFB reward (**Extended Data Fig. 5**) and trended toward lower prevalence of CF instructive signals (**Extended Data Fig. 8**), paralleling the lower motivation levels observed behaviorally (**Fig. 1**). Higher motivation with MFB reward may result from recruiting a wider and denser population of ascending DA axons—including both mesolimbic and nigrostriatal pathways^23,24^—compared to stimulating VTA somata alone. Similarly, operant behavior driven by CF stimulation elicited less prevalent GrC ramping signals, which again corresponded to lower behavioral motivation (**Extended Data Fig. 9**). These results thus distinguish GrC and CF signals from pure sensory encoding of expected stimuli.

Finally, our data along with many prior theories and studies suggest that GrC-CF-Purkinje cell microcircuits are likely especially well-suited to making precise computations of timing^55,56^, in this case reward timing. Thus, one possibility is that cerebellar timing estimates could help refine action-reward learning in nigrostriatal networks^57^. The high diversity of GrC information sources especially in lobule VI^58,59^ along with the immense GrC-Purkinje cell convergence ratio^60^ may enable cerebellar microcircuits to make exceedingly complex reward associations. Thus, cerebellar networks may be especially suited to extract obscure reward associations and their timing, especially early in learning when fast cerebellar plasticity processes may be most dominant^61^—an ability that could benefit a host of associative learning processes elsewhere in the brain.

Overall, demonstrating the existence of predictive and instructive encoding of DA rewards in the cerebellum, and their causal contributions to learning motivated behaviors, may more strongly link these circuits to broader brain reward networks and associated disorders, substantially broadening possible cerebellar contributions to reward-driven learning and behavior.

## Acknowledgements

We thank NIMH Section on Instrumentation for fabrication, the National Eye Institute Ocular Gene Therapy Core for viral packaging, and the National Eye Institute Building 49 Central Animal Facility staff for animal care and husbandry. We thank J.S. Diamond, L. Luo, and members of the Neocortex-cerebellum circuitry unit for helpful comments on the manuscript.

## Funding statement

BAF was supported by an NIH Center on Compulsive Behaviors Fellowship and Pre-Doctoral Fellow Intramural Training Award. This research was supported by the Intramural Research Program (ZIA NS009434-04) of the National Institutes of Health (NIH), National Institute of Neurological Disorders and Stroke (NINDS). The funders had no role in study design, data collection and analysis, decision to publish or preparation of the manuscript. The contributions of the NIH authors were made as part of their official duties as NIH federal employees, are in compliance with agency policy requirements, and are considered Works of the United States Government. However, the findings and conclusions presented in this paper are those of the author(s) and do not necessarily reflect the views of the NIH or the U.S. Department of Health and Human Services.

## Author contributions

BAF and MJW conceived of the study. BAF performed experiments. MJW, AO, LR, and PM assisted with stereotaxic surgeries. AO, SS, and LR assisted with histology. MJW, AO, SS, and ST assisted with behavioral studies. MJW assisted with microscopy. ST, SS, LD, and LR contributed to data processing. BAF and MJW analyzed data. MJW supervised the study.

## Competing interests statement

The authors declare no competing interests.

## Methods

### Experimental Model Details

#### Mice

Both male and female mice age 6-16 weeks were used and all mice were on mixed genetic background. Mice were maintained on a 12-hour light/dark cycle at 22°C and 40-60% relative humidity.

All GrC imaging experiments used Math1-cre (RRID:IMSR_JAX:011104) × Ai93 (TIGRE-LSL-TRE-GCaMP6f; RRID: IMSR_JAX:024103) × ztTA (R26-CAG-LSL-tTA; RRID: IMSR_JAX:012266) mice, which selectively express GCaMP6f in cerebellar GrCs.^62^ GrC imaging during optogenetic stimulation of VTA DA neurons used DAT-Flp (RRID: IMSR_JAX:035436) × Math1-cre × Ai93×ztTA mice, which express flippase in DA neurons.^30^

Imaging CF activity in Purkinje dendrites used viral induction of PkC-specific R-CaMP2 in Math1-cre×Ai93×ztTA or PCP2-Cre (RRID: IMSR_JAX:004146) × GCaMP8m (RRID: IMSR_JAX:037718), each with or without DAT-Flp. PCP2-Cre×GCaMP8m mice selectively express GCaMP8m^63^ in cerebellar PkCs.

Selectively inhibiting GrCs employed Math1-Cre×stGtACR1 (RRID:IMSR_JAX:037380) mice.

The following table summarizes cohort sizes and composition:

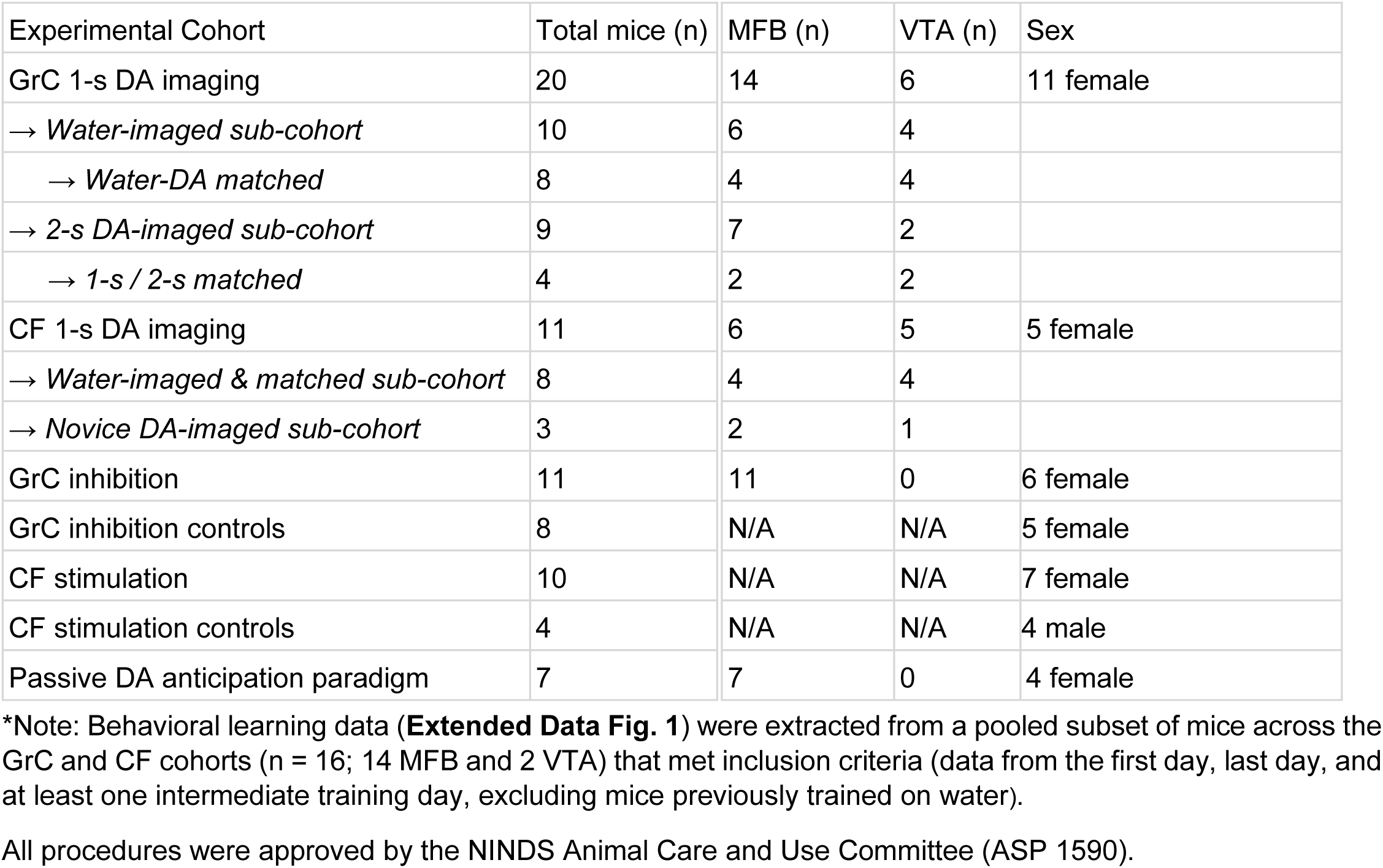

All procedures were approved by the NINDS Animal Care and Use Committee (ASP 1590).

### Method Details

#### Virus

In Math1-cre×Ai93×ztTA mice, PkC-specific promoter (L7-6)^64^ was used to drive expression of the red Ca^2+^ indicator R-CaMP2,^65^ which was packaged into AAV-L7-6-R-CaMP2. Plasmids were generated by VectorBuilder and packaged into AAV9 by the National Eye Institute Ocular Gene Therapy Core. Virus was injected at ∼10^12^ genomes/mL.

In DAT-Flp mice, Flp-dependent promoter (fDIO) drove expression of ChRimson^66^ in VTA DA neurons, transduced via AAV-Ef1a-fDIO-ChrimsonR-tdTomato. Virus was injected at ∼10^12^ genomes/mL.

For CF activation, AAV-CaMKIIa-ChRmine-mScarlet-Kv2.1-WPRE (RRID:Addgene_130991) was injected into the inferior olive to express opsin in CFs. Virus was injected at ∼10^12^ genomes/mL.

#### General Surgical Procedures

Surgery followed prior procedures^11^. Mice were anesthetized via isoflurane (∼2% in ∼1L/min of O_2_) and secured in a stereotaxic device (Kopf). We cleaned the scalp and removed hair using depilatory cream. We removed a ∼5x7 mm patch of skin, scraped the exposed skull surface, and used VetBond (3M) adhesive to seal the edges of the skin.

For DAT-Flp-expressing Math1-cre×Ai93×ztTA or PCP2-Cre×GCaMP8m mice, we infused 500 nL at 100 nL/min of AAV1-ef1a-fDIO-ChrimsonR-tdTomato (Addgene, RRID: Addgene_171027) into the VTA with a ∼25 μm diameter-tip glass capillary (coordinates: AP, -3.4 mm; ML, +0.5 mm; DV, -4.3 mm). Pipets were withdrawn after 5 minutes. A 5 mm length fiber optic cannula (Thorlabs CFML22L05 or CFMLC22L05, 0.22 NA, 0.2 mm diameter) was implanted (coordinates: AP, -3.4 mm; ML, +0.5 mm; DV, -4.1 mm) and secured with Metabond (Parkell UN1247).

For MFB experiments, a 5 mm length tungsten electrode (ProTech International, 8IMS303T3B01) dyed with DiD (Invitrogen, V22887) or FastGreen (Sigma Aldrich, F7252) dissolved in acetic acid was lowered into the MFB (coordinates: AP, -1.4 mm; ML, +1.2 mm; DV, - 4.8 mm) and secured with Metabond.

To express R-CaMP2 in PkCs, we drilled a craniotomy 0.3 mm right of the midline and centered on the post-lambda suture. We inserted a ∼25 μm diameter-tip glass capillary ∼300 um below the pial surface and injected 500 nL of AAV-L7-6-R-CaMP2 into the tissue and withdrew the pipet 5 minutes after injection.

To express ChRmine in the IO, the pipet was lowered at an azimuthal angle of 40 degrees, 0.5 mm left of midline, brought to the boundary between the cerebellum and the brainstem, and inserted 2.8 mm below the surface. 80nL was injected and then the pipet retracted 50 μm, to a maximum of 400nL volume and a minimum 2.6 mm insertion depth.

Experiments began after 2 weeks for virus expression.

### Histology

Mice were deeply anesthetized with isoflurane and transcardially perfused with 1x PBS followed by 10% formalin. The brain was removed, left overnight in 10% formalin, and then transferred to 30% sucrose solution in 1x PBS until it sank. We embedded brains in OCT Compound (Fisher HealthCare) prior to sectioning. Coronal or sagittal sections (50 μm) were collected using a Microm HM550 cryostat into well plates containing 1x PBS.

Sections were first washed 3 x 10 minutes with 0.1% PBS-T and blocked in 10% NGS in PBS-T.

For GrC GCaMP6f histology, we stained sagittal sections using rabbit anti-GFP (Abcam, AB290, 1:1500 dilution) for ∼48h, followed by 3 x 10 minutes washes with 0.1% PBS-T, followed by Alexa 488 goat anti-rabbit (Abcam, AB150077, 1:500 dilution) secondary antibody staining for ∼2 hours. PkC GCaMP8 histology used 1:2000 rabbit anti-GFP but was otherwise the same.

For DAT-Flp histology, we stained coronal midbrain sections using chicken-TH (Abcam, AB76442, 1:2000) for ∼48h at 4°C, followed by 3 x 10 min washes of 0.1% PBS-T, followed by Alexa 647 goat anti-chicken (Abcam, AB150083, 1:500 dilution) secondary antibody staining for ∼2 h.

We mounted sections with DAPI medium (Southern Biotech 0100-20) and imaged slides with a confocal microscope with 405, 488, 561, and 647 nm lasers (Zeiss LSM 510). ChRimson opsin (tdTomato) is visible under 561 nm illumination without staining. stGtACR1 opsin (mRuby) and ChRmine opsin were both visible without staining. MFB histology used no staining.

To analyze CF ChRmine histology, we segmented the image into GrC layer vs molecular layer. We created masks for all approximately 100 µm straight lines perpendicular to the folium in the molecular layer as putative CFs, and for all approximately 7 µm round structures in the GrC layer as putative mossy fiber boutons. CF:mossy fiber transduction specificity (212) was the ratio of counts normalized by the anatomical ratio of mossy fiber boutons to CFs in the mouse brain of ∼106 (400 GrCs/Purkinje cell × 4 mossy fiber synapses per GrC ÷ 15 GrC synapses per mossy fiber bouton).

#### Window and headplate implantation

Following previous approaches^11^, we drilled a ∼3.5 to 4 mm diameter cranial window centered over Lobule VI of the right vermis. We then used UV curing optical adhesive (Thorlabs NOA 81) to glue a 3 mm diameter glass coverslip onto a 3 mm outer diameter, 1 mm height, 2.7 mm inner diameter steel ring. We inserted the implant at a 15-20° angle counterclockwise from the sagittal axis and a 35-40° azimuthal angle from the coronal plane. We depressed the glass below the average depth of the underside of the skull at the window perimeter. We then sealed the steel ring to the skull using Metabond. Finally, we implanted a headplate (5 mm central opening and two extensions for fixation) parallel to the glass window and secured it with Metabond.

#### Behavioral Data Collection

Custom National Instruments cRIO software controlled and recorded the movement of a custom two-axis planar robotic manipulandum^26^. Through a single DAQ this device and software synchronously recorded: handle position; microscope frame counter TTLs; camera frame trigger TTLs; water solenoid and/or optogenetic laser and/or microwire stimulation TTLs. These signals were used to synchronize the behavior and imaging data post hoc and, in cases where dlc body tracking was performed, also the camera recordings.

#### Reach task behavioral Training

During all self-stimulation studies (VTA-DA optogenetics, MFB electrical activation, CF optogenetics, and 640 nm light-only control), mice were given ad libitum access to food and water. For water reward, mice were restricted to 0.4 mL of water per every 10 g of free feeding weight to maintain 80% of initial body weight. Mice were weighed daily to monitor for excess weight loss in addition to checking for signs of lethargy, coat deterioration, hunching, and general distress. Mice obtained water until satiety during behavioral training with additional water provided as needed.

All self-stimulation and push-for-water cohorts employed our custom two axis robotic manipulandum.^26^ Mice self-initiated trials by pushing the handle until one of two conditions were met: (1) position reached the 8 mm robotic “wall” or (2) movement was aborted (no motion for >100 ms at any distance >3 mm). Pushes >6 mm were deemed completed, while movements >3 and <6 mm were considered aborted. In all cases, at movement end, the motors locked the handle in its final position. Following a delay (either 1 or 2 s depending on the study), reward was delivered (rewarded trials) or withheld (aborted and reward omission trials). Following another 2-s delay, the motors automatically returned the handle to the animal over the next 2 s, after which the mouse was able to self-initiate the subsequent trial. Thus all trials shared the same temporal structure regardless of outcome, and reward was the only external sensory event between movement end and handle return (except: (1) MFB imaging studies, which generated no sensory stimulus at reward delivery; (2) GrC optogenetic inhibition studies during which the inhibition laser light was also a variably timed visual stimulus during the delay).

Training proceeded identically for all cohorts. Animals received one ∼18–20 minute training session/day. Because the task is self-paced, the total number of trials varied depending on the animal’s motivation. Training continued for 5–7 sessions before any imaging. Reward omission trials were only delivered in sessions 3+, and were pseudorandomized to ensure exactly 2 omission trials in each 10 trial block. No shaping steps were taken to establish the operant behavior or to encourage or discourage specific collateral body movements.

In push-for-VTA and push-for-CF stimulation sessions, light was delivered by a 640 nm laser (Coherent OBIS) triggered by the cRIO and routed through an optical fiber coupled to a fiber cannula (VTA) or the cranial window (CF and light control). In push-for-MFB activation sessions, the cRIO trigger drove an electrical current through the tungsten electrode targeting MFB. For water reward, a water droplet was delivered in front of the animal’s mouth via a gravity-fed reservoir through a solenoid valve.

For reward switching studies, we first trained mice on either push for stimulation or push for water without collecting images. After 5-7 training days, we imaged one or more fields and then switched to the other reward type for an additional 5-7 days. Finally, we returned to the original imaging field(s), aligning neurons when possible. 1-s-to-2-s delay switching followed the same regimen.

#### Two-photon microscopy

A custom two-photon microscope with mechanically articulating objective and both 920 nm (GCaMP) and remotely focused 1064 nm (R-CaMP2) excitation lasers was used for all imaging studies. GrC imaging used a 40x 0.8 NA Olympus objective while CF imaging for PCP2-Cre×GCaMP8m mice used a 16x 0.8 NA Nikon objective. Remote focusing used an Optotune electrically tunable lens to manually pull the laser focus up from the GrC layer (where the 920 nm was focused) into the Purkinje dendritic layer (typically ∼50-100 μm). Images were 512x512 pixels and frame rate 30 Hz. Chronic GrC imaging required angular alignment of the objective to the cranial window glass via red laser back reflection through an iris in the objective port; stage coordinates with respect to a “landmark” blood vessel; and finally manual alignment of the live image to the previous session’s mean image using fine motor and objective z-piezo positioning.

For imaging during VTA optogenetics, an additional 625 nm short pass filter and stacked emission filters reduced 640 nm laser incident on the PMTs. For the red channel, we also used post hoc temporal interpolation to remove remaining artifacts (**Extended Data Fig. 1** for complete description).

For imaging during CF optogenetics, additional dichroic mirrors and substitutions were used to combine the 920 nm and 640 nm lasers and deliver them into the tissue while excluding both from the emission path. The extended pulse duration (50 ms) precluded the temporal interpolation strategy used for VTA activation, but the much briefer total activation duration (also 50 ms) permitted simply discarding imaging data during the 640 nm laser pulse.

#### Granule cell optogenetic inhibition

For GrC inhibition studies, Math1-Cre×stGtACR1 mice were randomly divided into laser-ON (11 mice) or laser-OFF (8 mice) training cohorts and trained on the standard push-for-MFB activation task for 5-7 days. Laser-ON cohorts had an optical fiber positioned above the cranial window that delivered 5–10 mW of 488 nm laser light with a random onset latency and duration following handle displacement exceeding 6 mm, triggered by the cRIO hardware. Onset latency after threshold-crossing, 300±180 ms, and duration 370±84 ms (median±median absolute deviation [m.a.d.]). After training, laser-ON mice performed one “recovery” laser-OFF test behavioral session.

#### Climbing fiber optogenetic activation

For CF activation studies, mice were trained for 5-7 days to push for a single 50 ms pulse of 640 nm light (5– 10 mW) delivered either by optical fiber directly or coupled into a microscope objective positioned over the cranial window, delivered 1 s after movement completion. Control mice with identical cranial window surgeries but no CF opsin underwent identical training.

#### Two-photon microscopy data preprocessing

All data were first motion-corrected using sequential large-displacement rigid correction followed by small-displacement nonrigid correction both provided by NoRMCorre^67^. Videos were subsequently corrected for slow drifts by dividing out exponential fits to the frame-averaged fluorescence across the recording session. GrCs were sorted using cNMF^31^. CFs were sorted using PCA/ICA^32^. After automated sorting, we performed manual filter curation, splitting, and merging as needed. Final traces were obtained by back-applying the spatial filters to the movie and z-scoring each cell’s fluorescence trace relative to the mean and standard deviation of its entire recording and, for CFs, performing deconvolution and spike detection via thresholding and calculating continuous spike rates using a 150 ms moving average filter.

#### Passive dopamine anticipation paradigm

Modified versions of the custom NI software used for the reaching task were used to control the behavior. The FPGA was programmed to drive an amplifier (PUI Audio AMP1X1) with a 500-ms duration 35 to 40 kHz “chirp” at 50% duty cycle. The amplifier board drove an ultrasonic piezoelectric transmitter (PUI Audio UTR-1440K-TT- R) placed ∼20 cm from the mouse. This frequency range was selected to provide a salient auditory cue within the optimal murine hearing range but distinct from the 8 kHz resonant scanning galvanometers. The FPGA delivered the MFB stimulation reward as described above 1-s after tone offset, followed by a 2.25 s ITI. This highly predictable temporal structure was intentionally designed to maximize reward anticipation and ensure robust engagement of GrC temporal ramping in the absence of a required instrumental action.

#### Statistics and Reproducibility

No statistical method was used to predetermine sample size. Sample sizes reflected available animals and recordings meeting the inclusion and quality criteria described above; exact n values are reported in figure legends. Trials, cells, or sessions failing the specified behavioral, imaging, registration, or pose-tracking criteria were omitted from the relevant analyses. Animals in the stGtACR1 inhibition experiment were randomized to the specified experimental groups. Other experiments were not randomized because comparisons were performed within animals or between cohorts defined by the required genotype, surgery or stimulation paradigm. The Investigators were not blinded to allocation during experiments or outcome assessment. Results were assessed across independent animals and sessions. Unless otherwise specified, centers and error estimates are median ± standard error of the median (the standard deviation of 10,000 bootstrap medians). Two-group comparisons used two-sided Mann–Whitney U tests (unpaired) or Wilcoxon signed-rank tests (paired). Multiple comparisons used Bonferroni-adjusted p values unless otherwise stated.

### Data analysis

Data averaging throughout the text used several data alignment points: “movement endpoint” (or “movement” for short in some cases) referred to the last timepoint at which the handle displacement was <85% of its final position or, in cases where animals pushed very hard and exceeded the loose robotic “wall” at 8 mm displacement, the first time that the displacement exceeded 9 mm; “movement start” referred to the first time point following previous trial handle return that the speed (in 50 ms moving average windows) exceeded 55 mm / s; “reward” referred to the first timepoint at which either the water solenoid, or the MFB wire, or the VTA optogenetic laser trigger, received the first TTL rising edge.

### Body-movement analysis

DeepLabCut quality control was applied from −1 to 3 s relative to movement end. Trials were rejected if DLC lever position correlated <0.85 with encoder position, tracking likelihood (when available) was <0.9 continuously for >0.2 s, a coordinate gap exceeded 1 s, or >0.3 s total had speed >1,000 pixels/s. Remaining low-confidence (≤0.2 s) and missing-coordinate (≤1 s) gaps were linearly interpolated, and Euclidean frame-to-frame speeds were calculated for the lower jaw, nose, and ipsilateral/right fore- and hindpaws. Analyses used good reaches

≥6 mm with complete speed traces and ≥2 trials per session/body part; paired-condition analyses retained mice represented in both conditions. Traces were resampled to a common timebase; omission traces were additionally zero-phase filtered with a second-order Butterworth filter (normalized cutoff, 0.5). Variance-partitioning sessions required >10 trials complete across all available landmarks.

### Single-GrC generalization

Cross-context generalization was assessed using Benjamini–Hochberg correction of two-sided correlation p values (FDR q = 0.01). The stricter cutoff, obtained from the water–dopamine comparison (n = 1,098 tracked GrCs), corresponded to r ≥ 0.40 and was also applied to the 1-s/2-s comparison.

To compute GrC temporal scaling correlations, the 2-s delay period was uniformly downsampled by a factor of two to map it onto the identical temporal duration of the 1-s delay period.

CFs were classified as DA or water reward-responding if their trial-averaged spike rate in the [0.1, 0.2] s post-reward window exceeded 1.3 Hz.

### Variance Partitioning Analysis

Variance partitioning separated movement- and reward-expectation-related GrC activity. For each cell, rewarded- and omission-trial-averaged fluorescence traces were smoothed with a five-frame Gaussian (∼166 ms) and concatenated over 0–3 s (1-s delay) or 0–4 s (2-s delay) relative to movement.

Tracked jaw, nose, forepaw, and hindpaw speeds (the available subset for each session) were interpolated to imaging times, concatenated, z-scored, and reduced by PCA; the first two z-scored PCs were used as kinematic regressors. Two expectation regressors were defined by expected reward time T. From movement onset to T, they followed t/T and √(t/T). After reward, both decayed as exp(−t_post_/0.15). After omission, they decayed to zero over T as 1 − t_post_/T and 1 − (t_post_/T)², respectively. All regressors were causally convolved with an exponential GCaMP kernel (τ = 0.14 s).

Ordinary least-squares models were fit to each concatenated neural trace. The full model contained two kinematic and two expectation regressors. Unique kinematic and expectation contributions were ΔR²_kinematic_ = R²_full_ − R²_expectation_ and ΔR²_expectation_ = R²_full_ − R²_kinematic_, respectively; negative values were set to zero.

## Data availability statement

Data supporting this study are publicly available at Zenodo, DOI: 10.5281/zenodo.21621975. Source data are provided with this paper.

## Code availability statement

Custom analysis code is publicly available at Zenodo, DOI: 10.5281/zenodo.21621975.

---

**Extended Data Fig. 1.**
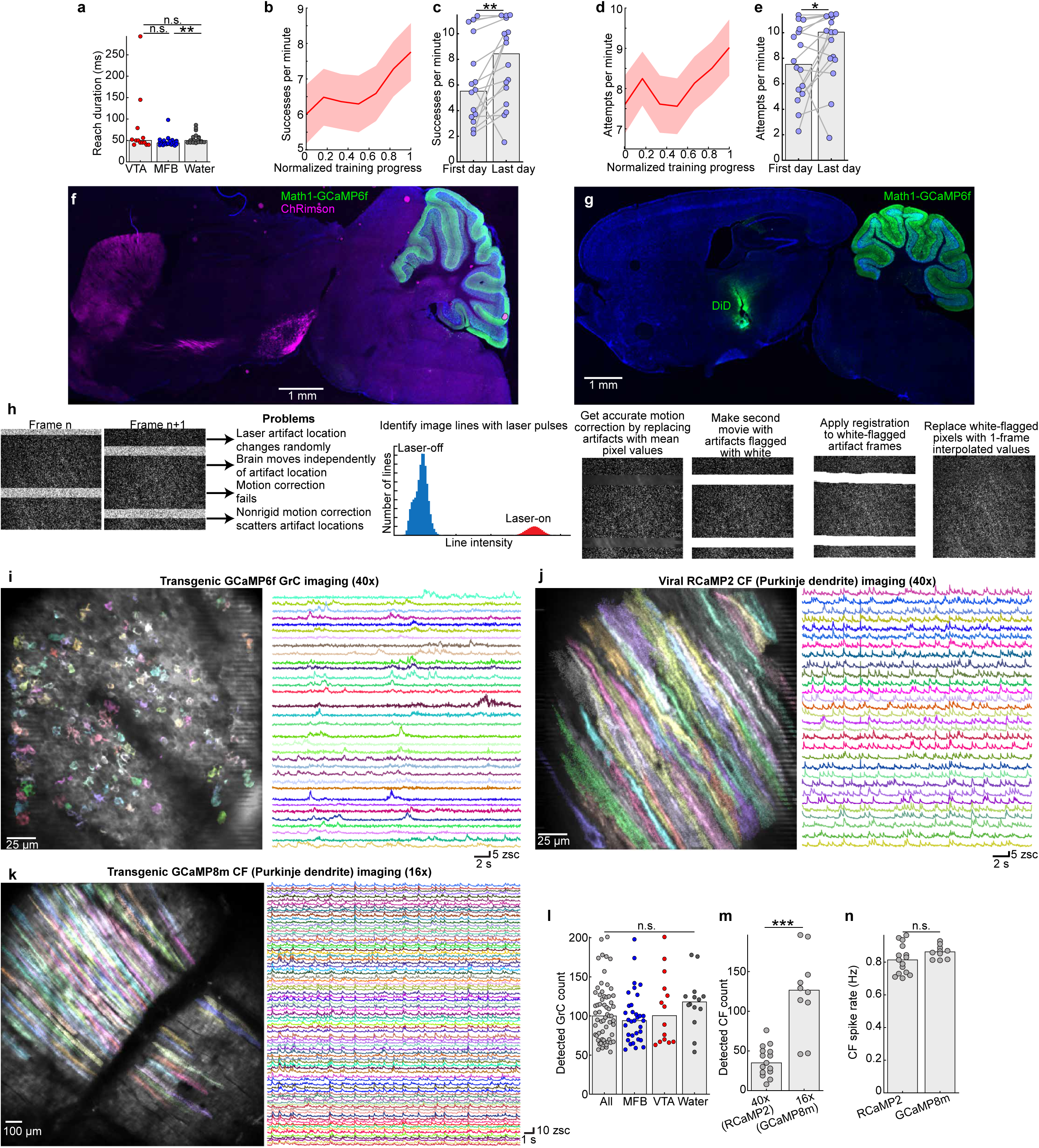

**Extended Data Fig. 2.**
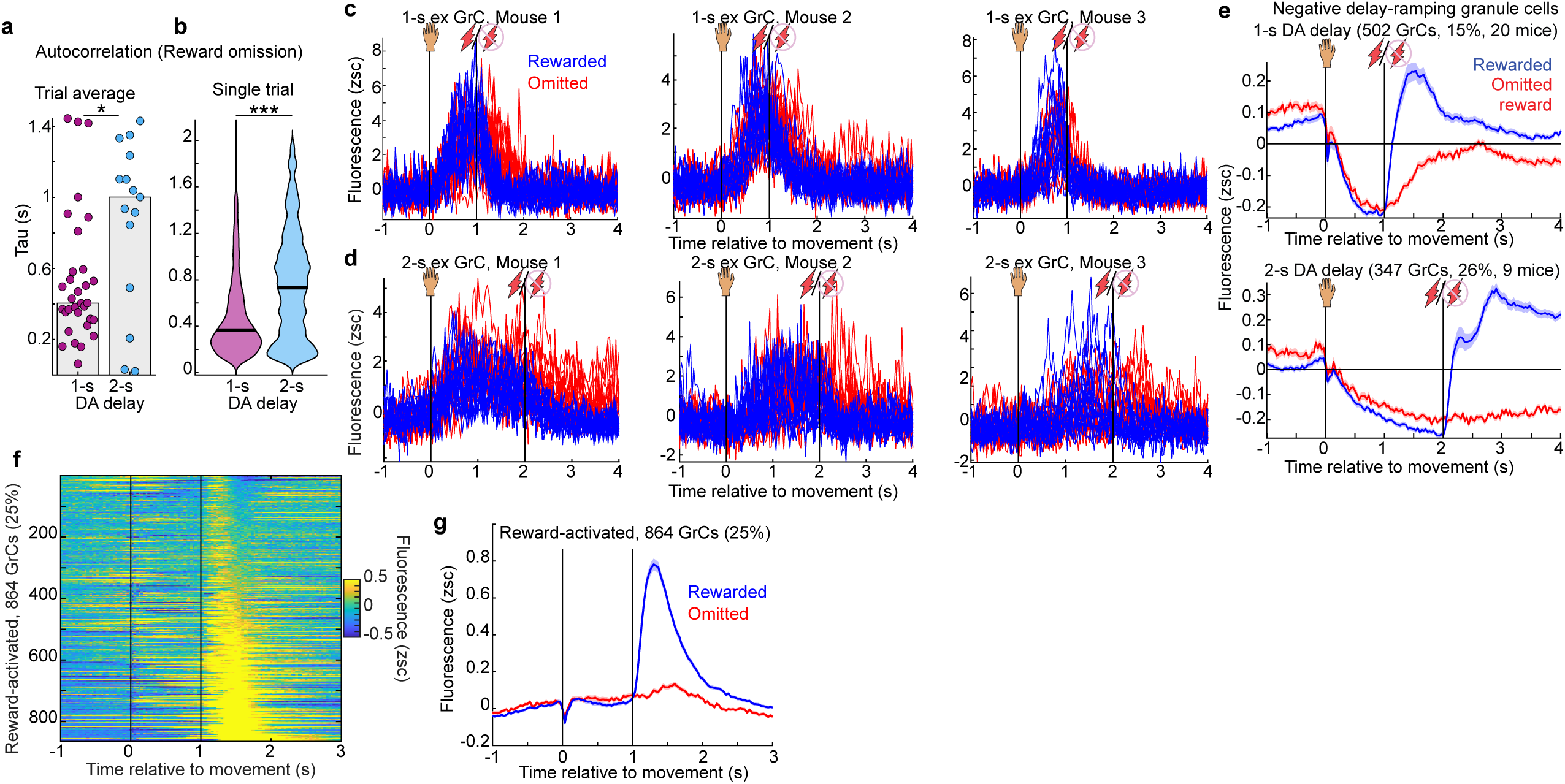

**Extended Data Fig. 3.**
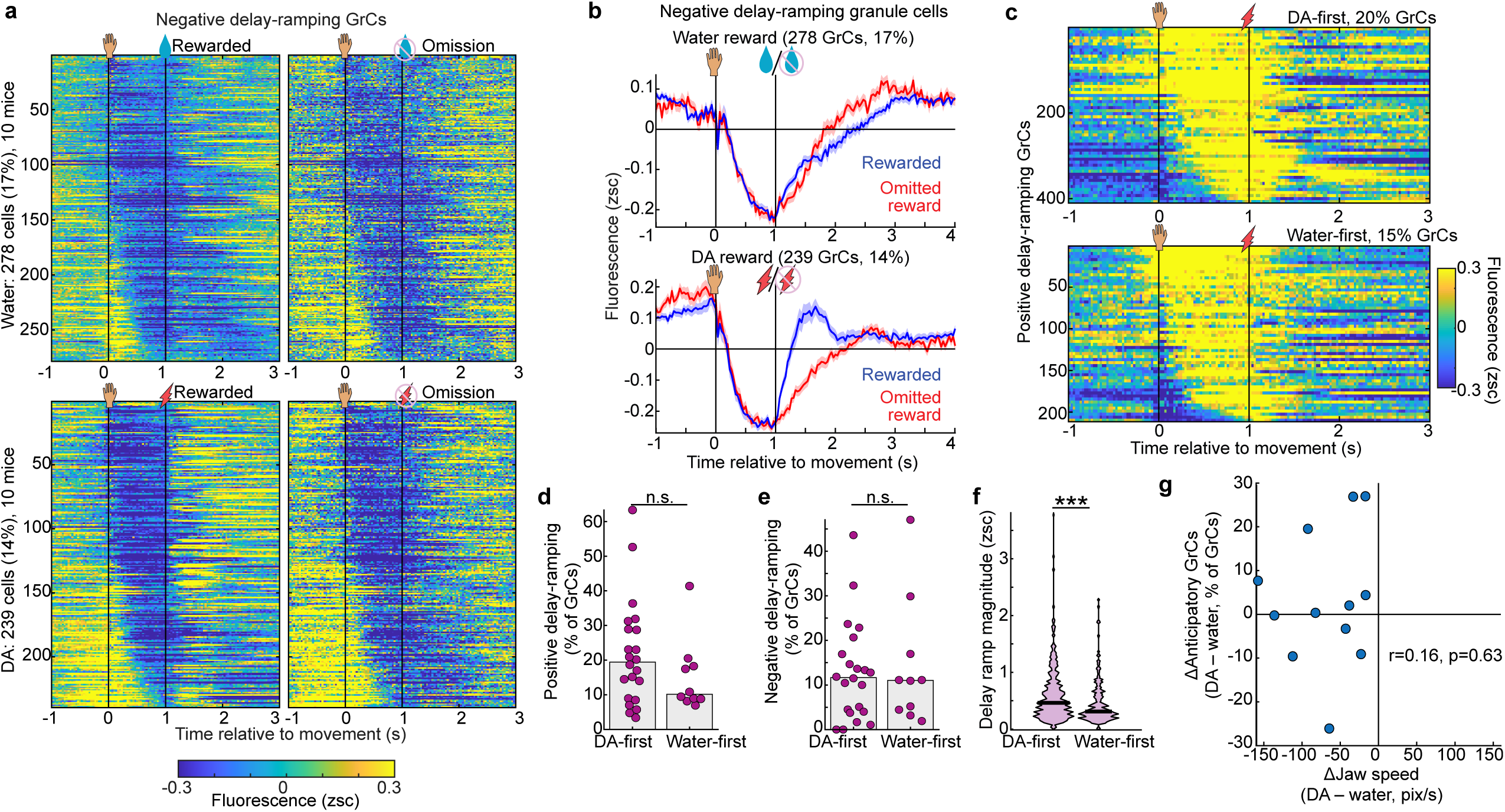

**Extended Data Fig. 4.**
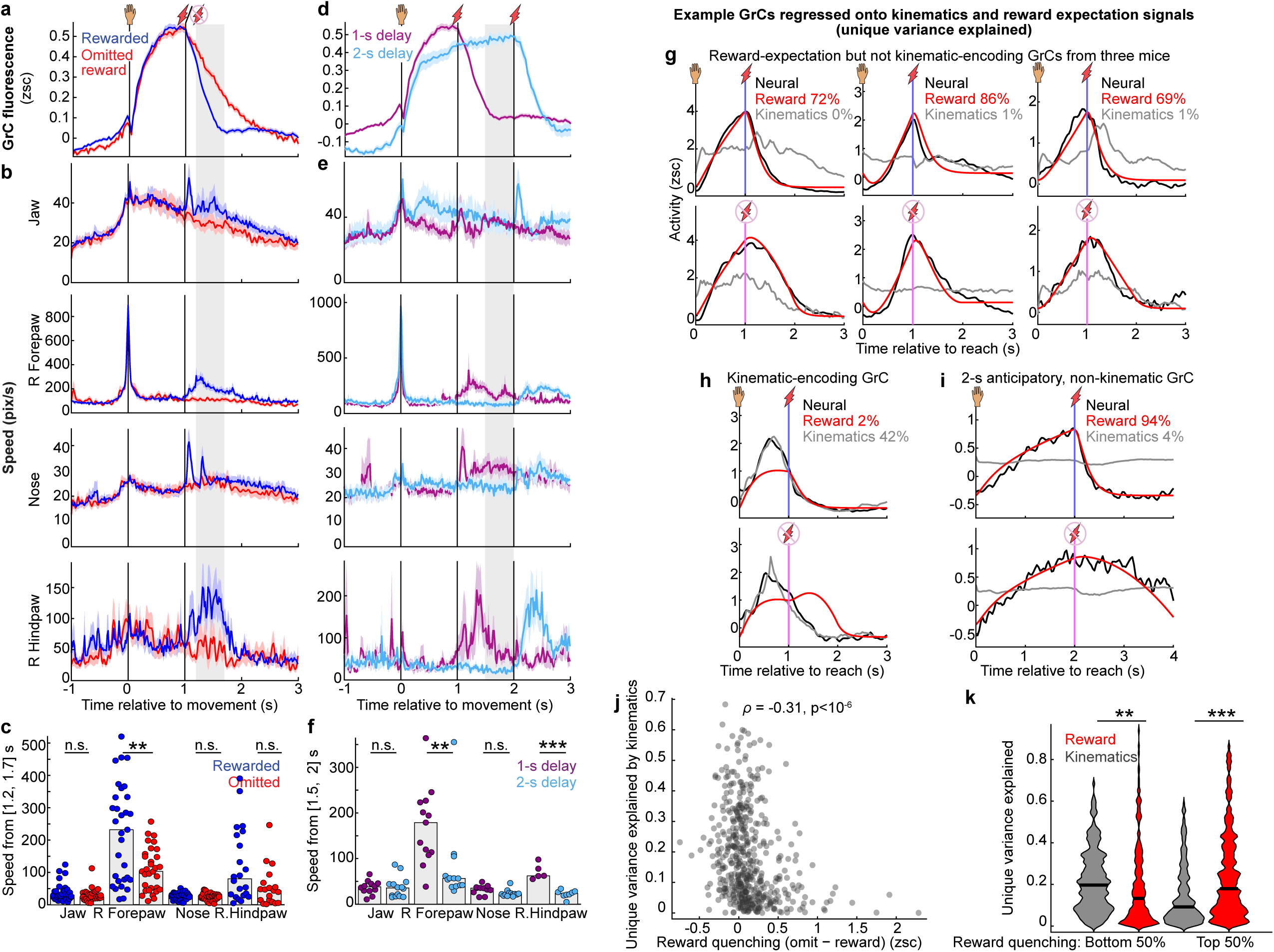

**Extended Data Fig. 5.**
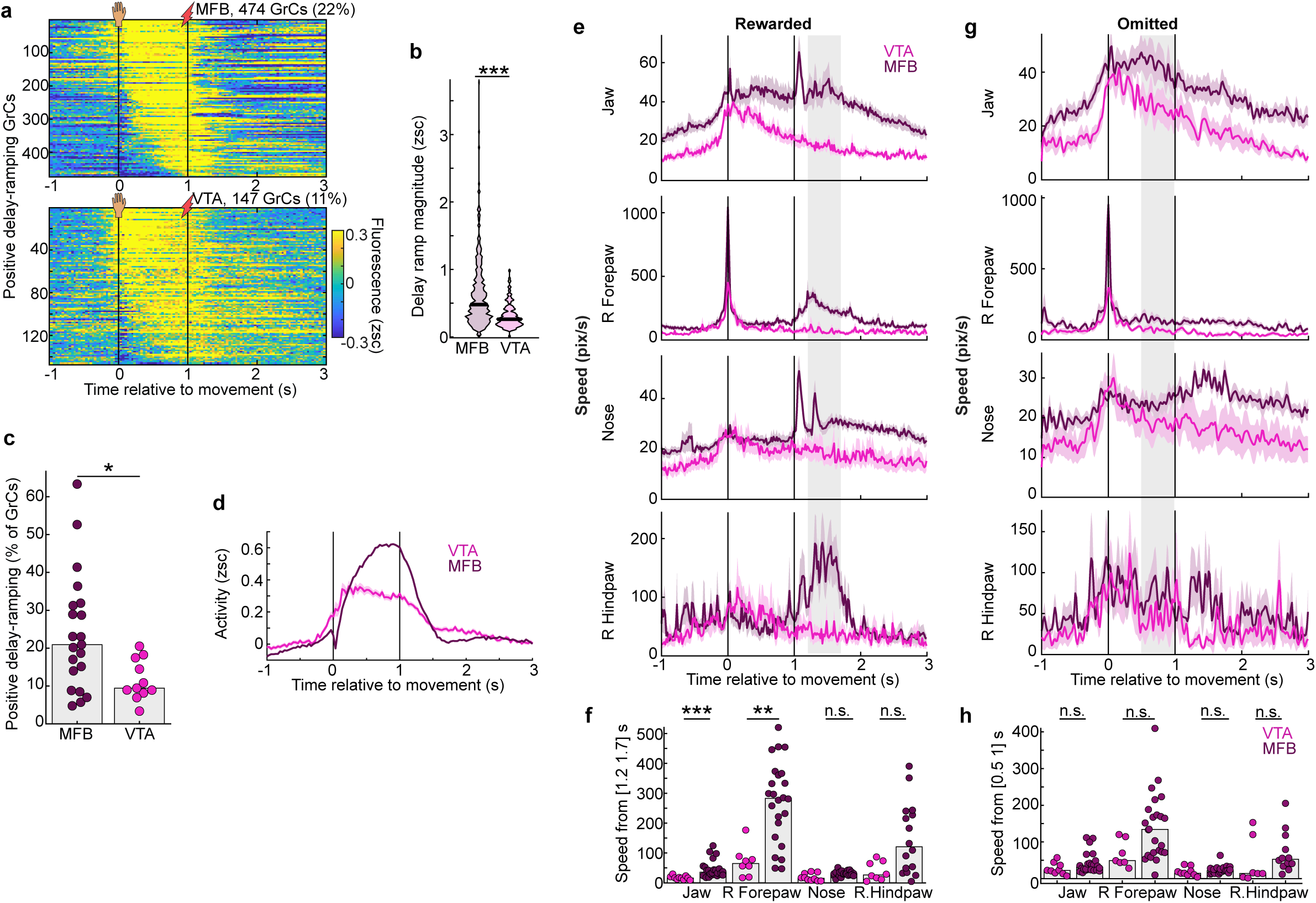

**Extended Data Fig. 6.**
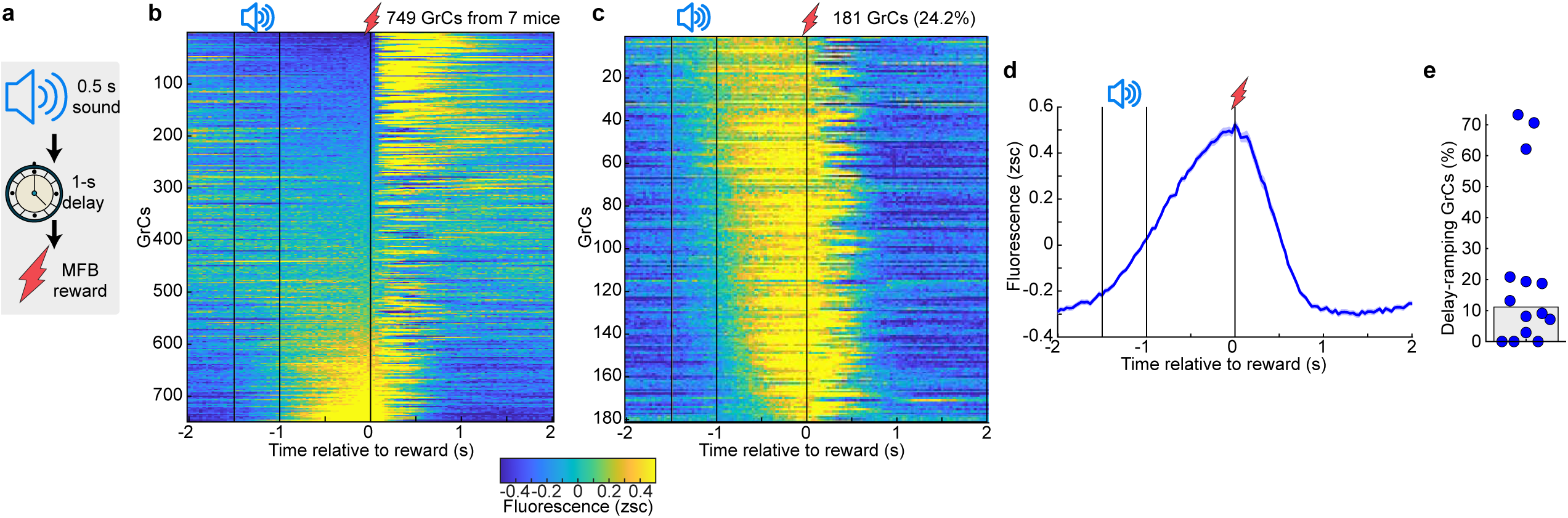

**Extended Data Fig. 7.**
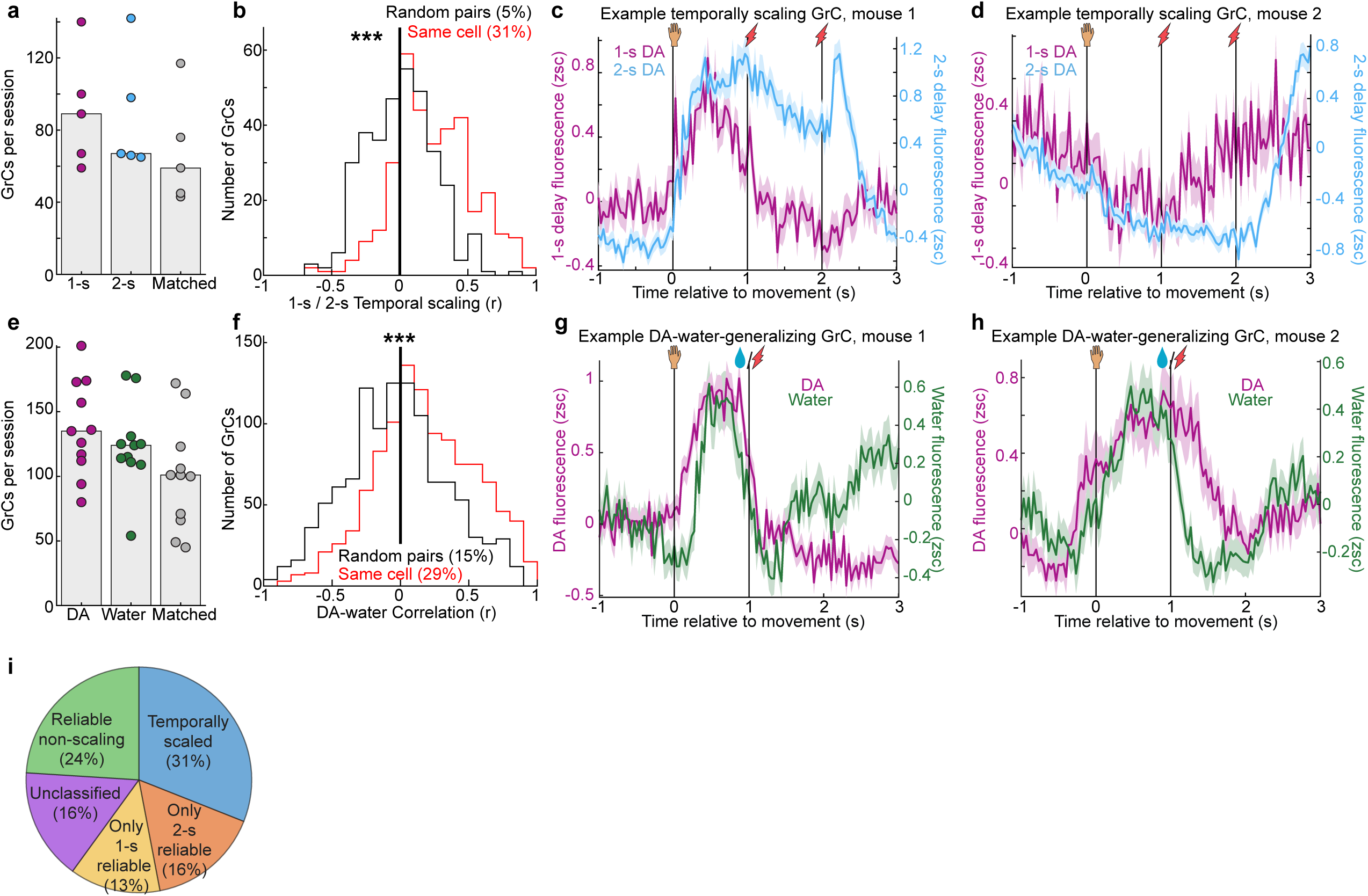

**Extended Data Fig. 8.**
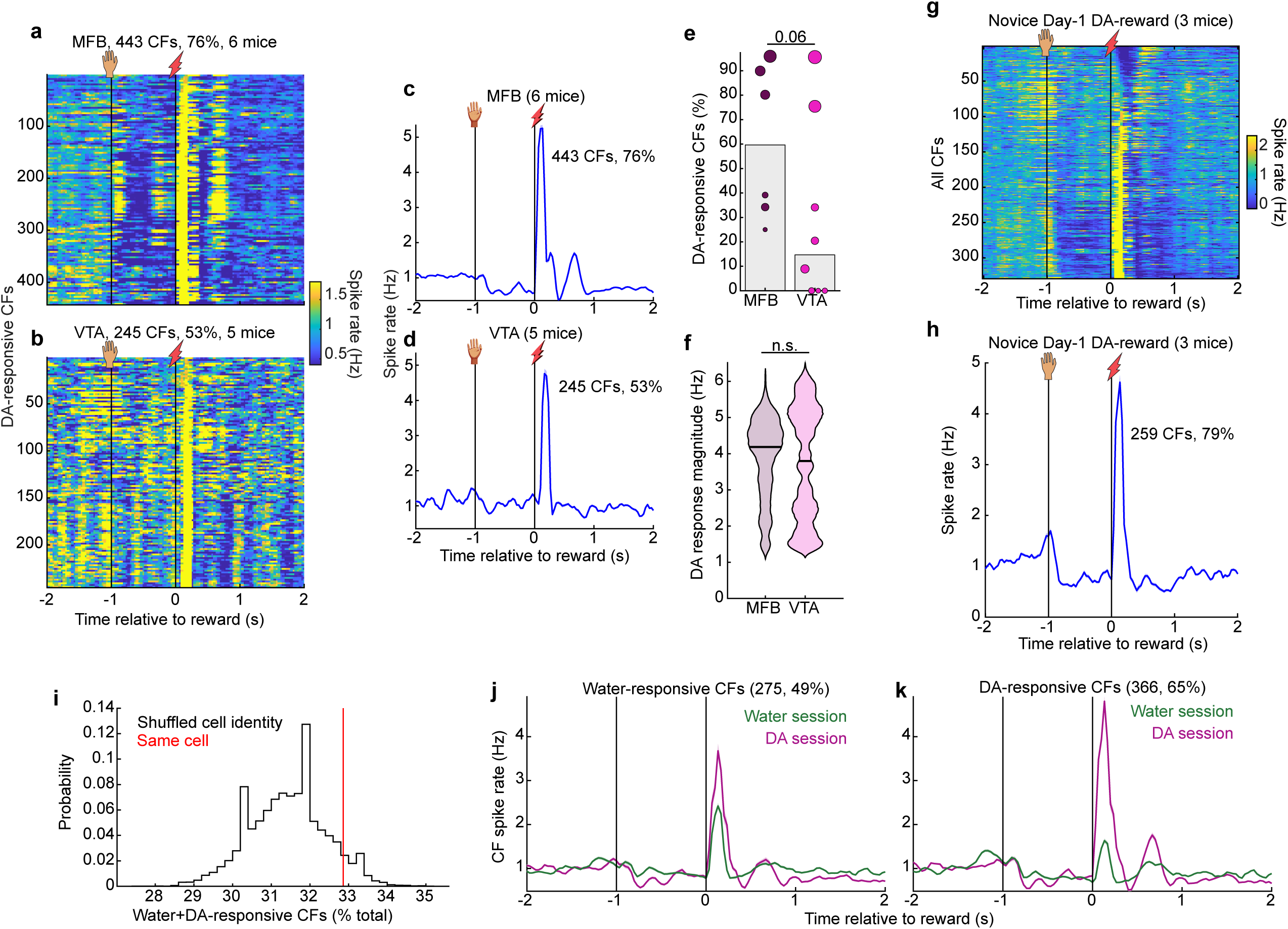

**Extended Data Fig. 9.**
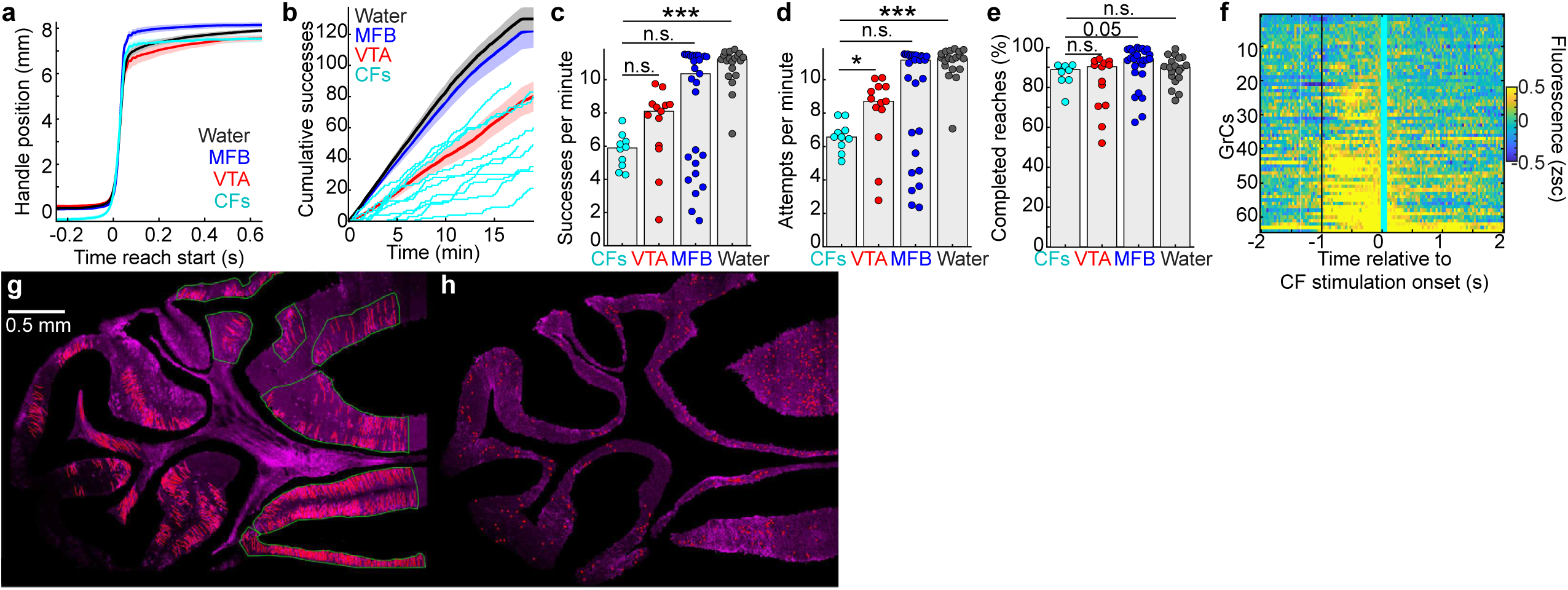

**Extended Data Fig. 10.**
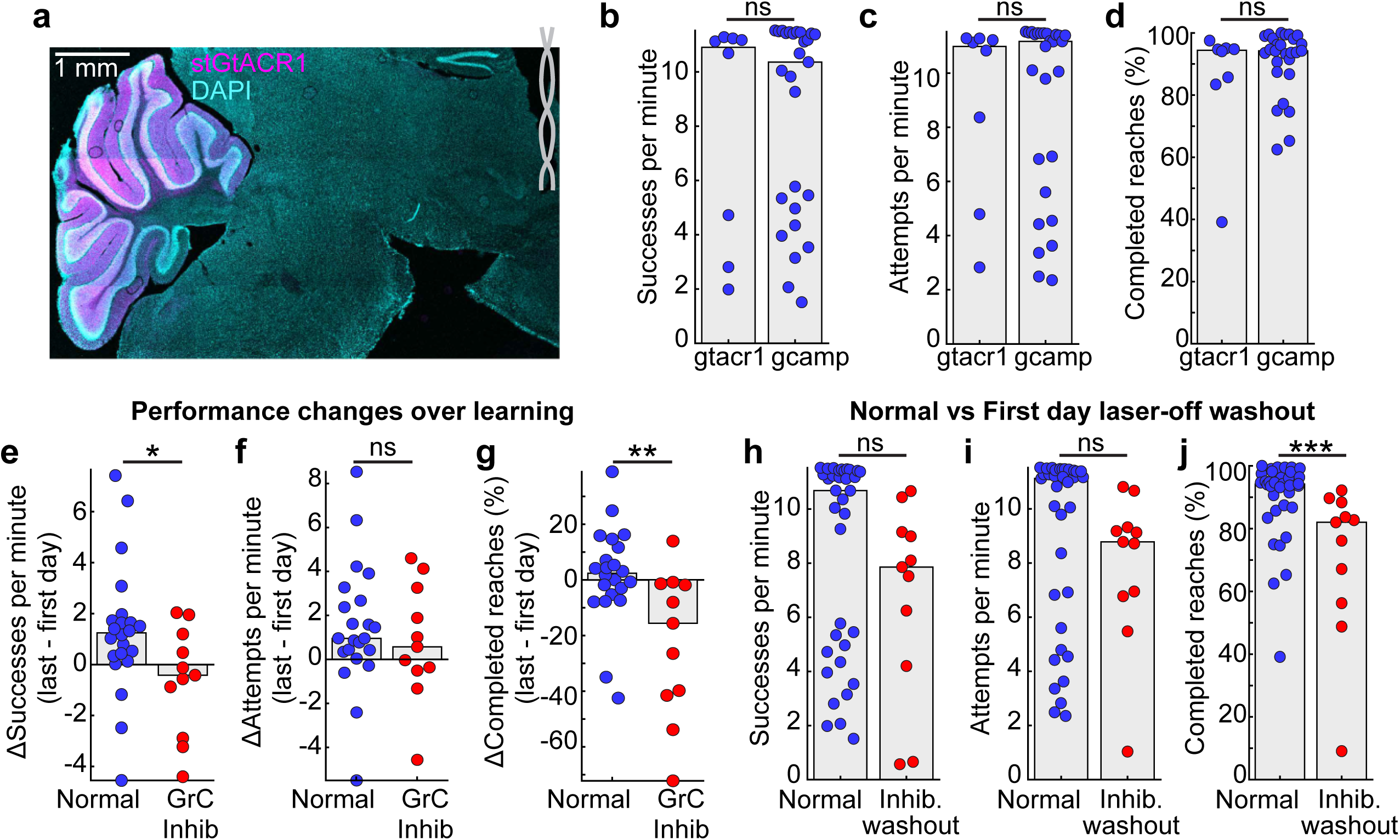

## Notes

https://doi.org/10.5281/zenodo.21621975

### Competing Interest Statement

The authors have declared no competing interest.

### Summary of Updates

Reduced to minimum size Figures 1-7 and ED Figures 1-10 for loading speed.

